# An hidden Markov model to estimate homozygous-by-descent probabilities associated with nested layers of ancestors

**DOI:** 10.1101/2021.05.25.445246

**Authors:** Tom Druet, Mathieu Gautier

## Abstract

Inbreeding results from the mating of related individuals and has negative consequences because it brings together deleterious variants in one individual. Genomic estimates of the inbreeding coefficients are preferred to pedigree-based estimators as they measure the realized inbreeding levels and they are more robust to pedigree errors. Several methods identifying homozygous-by-descent (HBD) segments with hidden Markov models (HMM) have been recently developed and are particularly valuable when the information is degraded or heterogeneous (e.g., low-fold sequencing, low marker density, heterogeneous genotype quality or variable marker spacing). We previously developed a multiple HBD class HMM where HBD segments are classified in different groups based on their length (e.g., recent versus old HBD segments) but we recently observed that for high inbreeding levels with many HBD segments, the estimated contributions might be biased towards more recent classes (i.e., associated with large HBD segments) although the overall estimated level of inbreeding remained unbiased. We herein propose a new model in which the HBD classification is modeled in successive nested levels with decreasing expected HBD segment lengths, the underlying exponential rates being directly related to the number of generations to the common ancestor. The non-HBD classes are now modeled as a mixture of HBD segments from later generations and shorter non-HBD segments (i.e., both with higher rates). The new model has improved statistical properties and performs better on simulated data compared to our previous version. We also show that the parameters of the model are easier to interpret and that the model is more robust to the choice of the number of classes. Overall, the new model results in an improved partitioning of inbreeding in different HBD classes and should be preferred.

## 1 Introduction

In diploid species, offspring of related individuals can carry at autosomal loci a pair of DNA segments originating from the same common ancestor. These stretches of contiguous loci where the two DNA copies are identical-by-descent (IBD) are referred to as homozygous-by-descent (HBD) or autozygous segments. The length of these HBD segments is inversely related to the size of the so-called inbreeding loop that connects the individual to its common ancestor, since multiple generations of recombination will tend to reduce the size of each transmitted DNA copy. The inbreeding level of an individual can be defined as the proportion of its genome that lies in HBD segments. Genomic data may allow to directly estimate this proportion to provide an estimator of the realized inbreeding coefficient (Leutenegger *et al*., 2003), whereas pedigree-based estimators, when available, can only provide expected values. Such estimates of inbreeding coefficients are highly valuable for the study of inbreeding depression and the management of livestock populations or those in conservation programs. In addition, detailed assessment of the distribution of HBD segments over the genomes can also be used in homozygosity mapping experiments (Abney *et al*., 2002; Leutenegger *et al*., 2006), to identify recessive alleles causing genetic defects or diseases, or for demographic inference purposes (Kirin *et al*., 2010; Ceballos *et al*., 2018).

In practice, HBD segments may be identified as runs-of-homozygosity (ROH) that correspond to long stretches of homozygous genotypes (Broman and Weber, 1999; McQuillan *et al*., 2008). Such ROH can be empirically detected with rule-based approaches requiring the definition of parameters such as window size, minimum ROH length, marker density, maximum allowed spacing between successive markers and number of missing or heterozygous genotypes (Purcell *et al*., 2007). More formally, likelihood-based ROH approaches allow to compare the likelihoods of segments to be allozygous versus autozygous regions based on marker allele frequencies and the genotyping error probabilities (Pemberton *et al*., 2012; Wang *et al*., 2009). These approaches still require the prior definition of fixed-length windows to scan the genome for ROH segments. Alternatively, several authors developed fully probabilistic approaches based on hidden Markov models (HMM) (Leutenegger *et al*., 2003; Narasimhan *et al*., 2016; Vieira *et al*., 2016; Druet and Gautier, 2017). As likelihood-based approaches, they rely on genotype frequencies and genotyping error probabilities, but in addition, they take into account inter-marker genetic distances. Moreover, they do not require prior selection of some window size as HBD estimations are integrated over all possible window lengths. Uncertainty in genotype calling, as for low-fold sequencing data, can also be integrated over (Vieira *et al*., 2016; Druet and Gautier, 2017). These two later characteristics thus make HMM methods particularly valuable for the analyses of data set with low marker density and/or heterogeneous genotype quality and/or heterogeneous marker spacing. For instance, they are the method of choice to work with ancient DNA (e.g., Renaud *et al*., 2019; Ringbauer *et al*., 2021), where genotype quality is particularly poor, and several HMM have been developed in the field. Similarly, they are particularly well suited to work with exome sequencing data (Magi *et al*., 2014), low density marker array (e.g., Solé *et al*., 2017; Druet *et al*., 2020) or with low-fold sequencing data (Vieira *et al*., 2016). Overall, fewer parameters need to be defined when using these tools.

In HMM based approaches, the length of HBD segments is generally assumed to be exponentially distributed. Modeling a single exponential distribution amounts to assume that all the autozygosity is associated to ancestors present in the same past generations. For complex population histories, this assumption may be too restrictive and Druet and Gautier (2017) proposed to use a mixture of exponential distributions to model HBD segment classes of different expected lengths, under a similar HMM framework. Such classification is actually an HMM counterpart to methods that were developed to automatically classify the identified ROH based on their observed length (Pemberton *et al*., 2012; Szpiech *et al*., 2017). With HMM, HBD classes can be viewed as group of ancestors present in different past generations. This model better accounts for complex demographic histories in which different ancestors from many different past generations may contribute to autozygosity. We showed that it improves the fit of individual genetic data and provides more accurate estimations of autozygosity levels. For instance, a single HBD class model might underestimate autozygosity when multiple generations contribute to it, and also tend to regress length of HBD segment towards intermediate values, cutting in particular the longest segments into shorter pieces (e.g., Solé *et al*., 2017). An accurate estimation of HBD segment length distribution may however be critical to estimate the number of generations to the common ancestors. Likewise, the multiple HBD-class model provides insights into the past demographic history of populations by estimating the relative contributions of past generations to contemporary inbreeding levels (Druet and Gautier, 2017).

The properties of the multiple HBD-class model have already been studied in detail in some of our previous works and most particularly its robustness to i) HBD segment length (age of the ancestors); ii) marker density; iii) allele frequency spectrum; iv) sequencing depth; v) genotyping error; and vi) variable recombination rate (e.g., Solé *et al*., 2017; Druet *et al*., 2020). We also investigated aspects related to model selection, including the number of classes and their rates, in simulated (Druet and Gautier, 2017) and real data sets (Solé *et al*., 2017). The behaviour of these models were also evaluated when multiple-HBD classes contributed to autozygosity, including comparisons when the underlying ancestors were separated by a few generations. A detailed discussion on these aspects is for instance available in Druet and Gautier (2017) and in the vignette from the RZooRoH package (Bertrand *et al*., 2019). Importantly, we showed that the model was robust to background Linkage Disequilibrium (LD) that was well captured by the most ancient HBD classes (i.e., HBD segments) and thus, as desired, did not influence estimates of recent inbreeding levels (Druet and Gautier, 2017; Solé *et al*., 2017). From a practical point of view, this made LD pruning of the analyzed data set unnecessary. In addition, the multiple-HBD class model has been compared to other methods, including rule-based ROH, likelihood-based ROH and the single HBD class HMM (Druet and Gautier, 2017; Solé *et al*., 2017; Alemu *et al*., 2021).

We recently observed that when the contribution of recent ancestors is extremely high, the multiple HBD classes model in its initial definition (as of Druet and Gautier, 2017) tended to underestimate the age of HBD segments by shifting HBD partitioning towards more recent classes (Druet *et al*., 2020), although the overall estimated levels of inbreeding remained unbiased. To solve this issue we herein propose a modified model in which the HBD classification is modeled in successive nested levels, each level corresponding to a single HBD class model with a distinct rate. As a result, the non-HBD classes are now modeled as a mixture of HBD segments from later generations and shorter non-HBD segments (i.e., both from subsequent levels with higher rates). We carried out a detailed simulation study to show that the upgraded model had better statistical properties and performed better compared to our previous version. We also show that the parameters of the model are easier to interpret and that the model is more robust to the choice of the number of classes (e.g., the autozygosity partitioning remains more similar when additional classes are added). We also provide an illustration on genotyping data from European Bison that we previously analyzed with the original model (Druet *et al*., 2020).

## 2 Models

### 2.1 Previous models

#### 2.1.1 Single HBD-class model (1R model)

Leutenegger *et al*. (2003) proposed to describe the genome of an individual as a mosaic of HBD and non-HBD segments with a HMM. In that model, the length of HBD segments inherited without recombination from a common ancestor is exponentially distributed with a rate *R*. This rate *R* is related to the number of generations of recombination along both paths connecting each of the two individual DNA copies (haplotype) to their common ancestor, and their frequency is a direct function of the mixing coefficient *ρ*. The HBD and non-HBD segments are not directly observed but their distribution can be inferred using genotype data available for a set of markers. In that case, the model can be represented as an HMM with two hidden states (state 1 = “HBD” and state 2 = “non-HBD”) with the following transition probabilities between two consecutive markers *m* and *m* + 1:

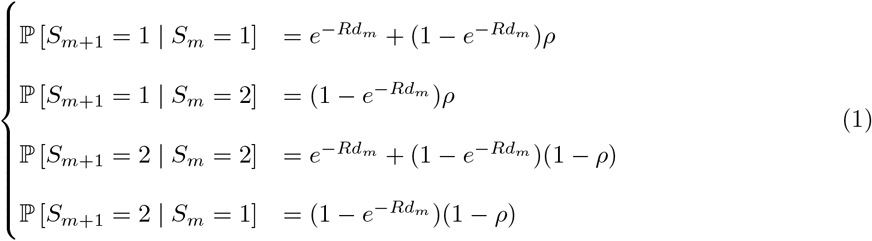

where *S*_*m*_ is the state at position *m, d*_*m*_ is the genetic distance in Morgans between markers *m* and *m* + 1. The term 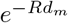 represents the probability that there is no recombination on both genealogical paths between two consecutive markers *m* and *m* + 1 (i.e., the HBD status remains the same). We use the term ‘coancestry changes’ to refer to the presence of at least one recombination on these paths as in Leutenegger *et al*. (2003), and *R* will be called the rate of coancestry change accordingly. In this HMM, the equilibrium HBD probability is *ρ*, which has been shown to be an unbiased estimator of the inbreeding coefficient defined as the proportion of genome HBD (Leutenegger *et al*., 2003). Note that the inbreeding coefficient may also be derived from the estimated posterior HBD probability at each marker (see eq. 21 below) leading to slightly different but highly correlated estimations.

It should be noticed that the expected length of HBD segments, that we define here and in the remainder of our work in a strict sense (i.e., without any coancestry change), is equal to 1*/R*. Yet, in individual genomes, some HBD segments may actually be neighboring. For instance, in the case of a marriage between cousins a tract of IBD markers may consist of two consecutive HBD segments inherited from the grand-father and the grand-mother. More precisely, under the above 1R model, the number of consecutive HBD segments actually follows a geometric distribution with parameter 1 *− ρ*, the probability of entering a non-HBD segment after a coancestry change. As a result, the expected length of tracts of IBD markers that may include one or several coancestry changes is equal to 1*/R*(1 *− ρ*) as noticed by Leutenegger *et al*. (2003).

The emission probabilities are the probabilities to observe the marker genotypes conditionally on the underlying state. For non-HBD and HBD states, these emission probabilities are a function of expected genotype frequencies in non-HBD and HBD segments, respectively (Crow *et al*., 1970; Broman and Weber, 1999; Leutenegger *et al*., 2003). For the HBD state:

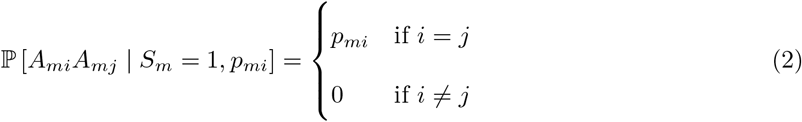

where *A*_*mi*_ and *A*_*mj*_ are the two alleles observed at marker *m, i* and *j* representing the allele numbers, *p*_*mi*_ is the frequency of allele *i* at marker *m*. Ideally, these allele frequencies should be estimated from individuals in a reference population but they are generally computed from the sampled individuals. For the non-HBD state:

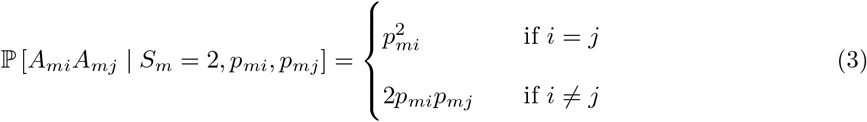

The expected frequencies in non-HBD segments (eqn. 3) correspond to Hardy-Weinberg proportions. These emission probabilities are similar to probabilities used in maximum likelihood estimators of the inbreeding coefficient (e.g., Weir *et al*., 2006). As a result, when markers are considered independent (i.e., probability of coancestry change equal to 1), both approaches lead to very similar estimates (see Alemu *et al*., 2021). The extension of these emission probabilities to incorporate genotyping error or mutation probability is straightforward (see Broman and Weber, 1999; Leutenegger *et al*., 2003; Druet and Gautier, 2017). Similarly, the emission probabilities can also be modified to handle next-generation sequencing data (e.g., genotype likelihoods) allowing efficient analysis of shallow sequencing or GBS data (see Vieira *et al*., 2016; Narasimhan *et al*., 2016; Druet and Gautier, 2017).

#### 2.1.2 Models with multiple HBD classes (KR and MixKR models)

In the single HBD class model, all HBD segments have the same expected length defined by the rate parameter *R*. Hence, ancestors contributing to HBD segments are assumed to have been present approximately in the same past generations. To model the contribution of different groups of ancestors to autozygosity (i.e., account for the difference in HBD segment lengths originating from ancestors living in different past generations), we introduced models with multiple HBD classes (Druet and Gautier, 2017). In these new models, each class correspond to a distinct state, with states 1 to *K −* 1 for HBD segments originating from groups of ancestors living in different past generations and a *K*th state for non-HBD positions. For each HBD class *c* (*c* = 1, …, *K −* 1), HBD segment lengths are assumed exponentially distributed with rate *R*_*c*_. The non-HBD state corresponds to positions that do not trace back to the same haplotype from a common ancestor up to the most remote HBD class, and has it own rate *R*_*K*_. The transition probabilities from state *b* at marker *m* to state *a* at marker *m* + 1 are:

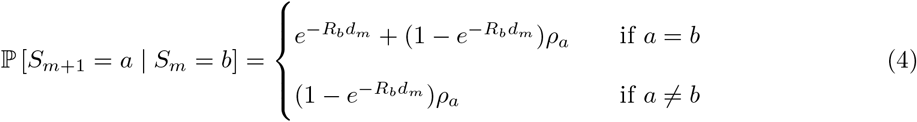

where *ρ*_*c*_ is the mixing coefficient associated with class *c*.

We previously called these multiple rate models, ‘KR’ models (e.g., 1R model corresponding to the single HBD-class model) where the number *K* refers to the total number of states (i.e., pertaining to *K −* 1 HBD classes and 1 non-HBD class). We proposed to estimate for each individual either the *K* different rates *R*_*c*_ or to set these rates to pre-defined values (so-called MixKR model) (Druet and Gautier, 2017). In the latter case, the rate for the non-HBD class was set equal to the most remote HBD class (i.e., *R*_*K*_ = *R*_*K−*1_). In practice, the MixKR modeling facilitates comparisons across different individuals and in the present work we only consider MixKR models. More importantly, the estimated *ρ*_*c*_ mixing coefficients associated to each HBD class *c* in KR models (with *K >* 2) can no longer be interpreted as inbreeding coefficients as in the single HBD class model. Indeed, although they correspond to the initial HMM state probabilities, the *ρ*_*c*_ values do not correspond to the marginal equilibrium proportions of genomes belonging to each HBD class *c* because these proportions also depends on the rates *R*_*c*_ that now differ between classes. Nevertheless, several measures related to individual inbreeding coefficients can be obtained from KR models as i) the genome-wide estimate of the realized individual inbreeding level 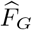, corresponding to the proportion of the genome in HBD classes; ii) the inbreeding level 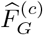 associated with HBD class *c* defined as the proportion of the genome belonging to class *c*; and iii) the posterior HBD probability *φ*_*m*_ corresponding to the probability that a locus *m* lies in a HBD segment (Druet and Gautier, 2017).

In addition to the loss of interpretability of mixing coefficients, we previously showed that the MixKR model tended to assign HBD segments to more recent classes (i.e., with smaller *R*) when the overall inbreeding level of individuals was high (Druet *et al*., 2020). Although the 1R model remained limited in its range of applications (because it models a single class of ancestors, see above), it provided both an unbiased estimate of *R* and an estimate of *ρ* that could be interpreted as an inbreeding coefficient.

### 2.2 New model: the nested 1R model

Here were propose a modified multiple HBD classes model that preserves the desirable properties of the 1R model and allows for the contribution of multiple groups of ancestors to autozygosity (as in the MixKR model). As illustrated in Figure 1, we sequentially model multiple layers of ancestors (from the most recent to the oldest), each contributing to a distinct HBD class. More precisely, a 1R model is first used to describe the genome of an individual as a mosaic of HBD segments associated with the most recent layer of ancestors (first group of ancestors) and non-HBD segments (i.e., relative to these ancestors). Although these positions would be non-HBD with respect to this first layer, they could be inherited HBD from more remote ancestors. Therefore, we propose to model in turn the non-HBD positions in the first layer as a mosaic of HBD and non-HBD segments associated with a second layer of ancestors (see Figure 1). This would be achieved by fitting a second 1R model, nested in the first one, with different parameters, *ρ*_2_ and *R*_2_ (with *R*_2_ *> R*_1_). This approach can be repeated for several layers of ancestors (Figure 1).

**Figure 1.**
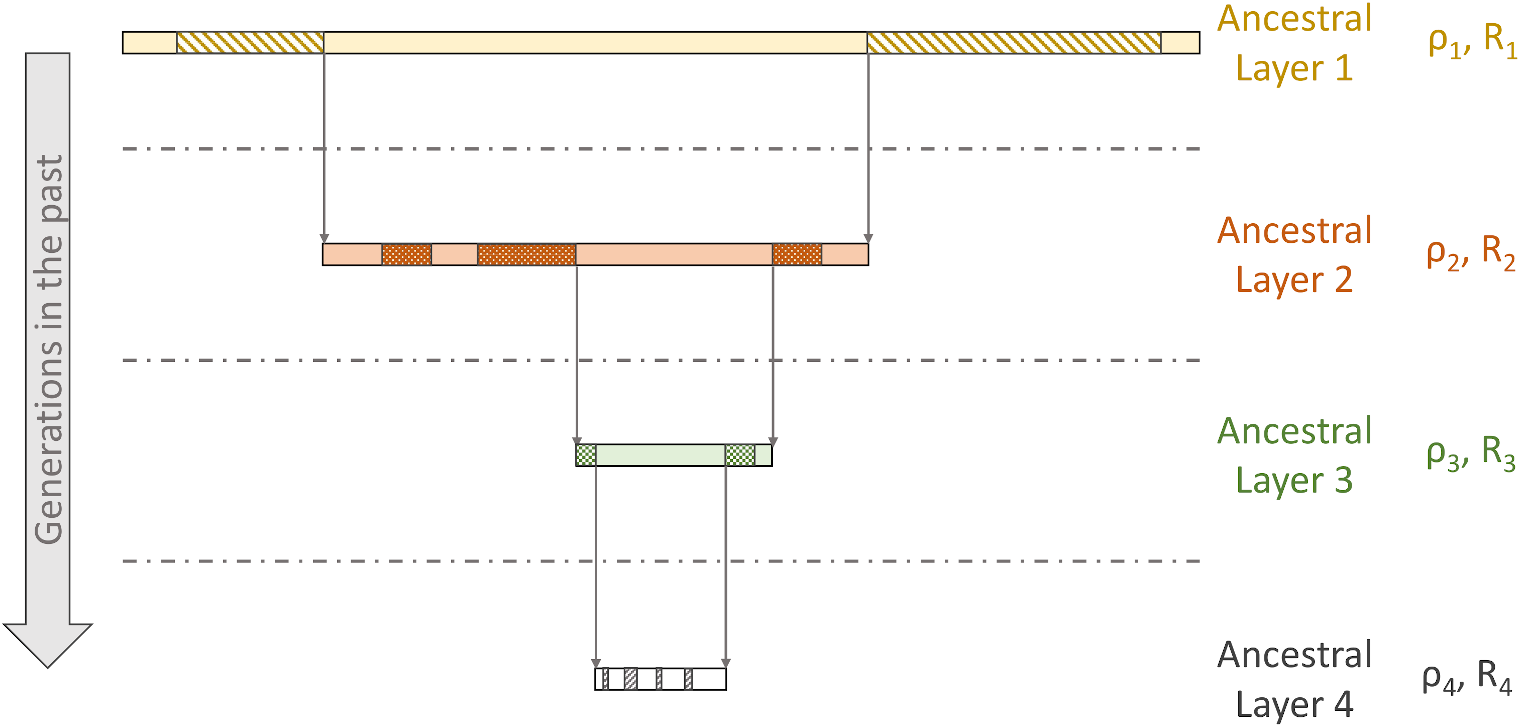
Graphical illustration of the Nested 1R model. Four layers of ancestors are represented. In each layer, the genome is represented as a mosaic of HBD and non-HBD segments with a 1R model with specific parameters *ρ*_*c*_ and *R*_*c*_. Regions with motives correspond to HBD segments.

Each layer *c* is thus described as a mosaic of HBD and non-HBD segments, labelled as HBD_*c*_ and non-HBD_*c*_ states (we use subscript *c* as layers match with HBD classes). The non-HBD class in layer *c* would be a mixture of HBD classes in subsequent layers and the non-HBD class in the last layer *L* (the total number of layers *L* = *K −* 1, where *K* is the number of hidden states). We assume that emission probabilities in HBD classes are the same in each layer, and identical to those used in the 1R model (eqn 2). Note that emission probabilities could be made layer dependent, e.g., to account for more generations of mutation or changes in allele frequencies through generations. Similarly, the emission probabilities for the non-HBD class in the last layer *L* matches those used in the 1R model (eq 3). However for non-HBD positions in layer *c* = 1 to *c* = *L −* 1, the emission probabilities now also depend on the mixing coefficients *ρ*_*c*_ through the proportion 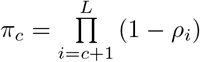 of positions expected to ultimately lie in a non-HBD segment at the oldest layer *L* (i.e., not mapping to an HBD segment in any successive layers *c*′ *> c*) as:

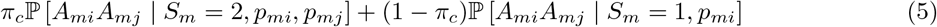

where ℙ [*A*_*mi*_*A*_*mj*_ | *S*_*m*_ = 2, *p*_*mi*_, *p*_*mj*_] and P [*A*_*mi*_*A*_*mj*_ | *S*_*m*_ = 1, *p*_*mi*_] are emission probabilities from the 1R model (eqns. 2 and 3).

As the parameters *ρ*_*c*_ for the different classes are required to obtain these emission probabilities, the implementation of this model is not trivial. A more convenient way to specify the Nested 1R model is to define *L* HBD states (one per layer) and a single non-HBD class associated to the *L*th layer. This results in a parameterization very similar to a MixKR model with a number of hidden states *K* = *L* + 1 (Druet and Gautier, 2017) but with a modified transition probabilities matrix **T**^**m**^ between consecutive markers *m* and *m* + 1. More precisely, in the MixKR model, **T**^**m**^ can be decomposed in three parts i) a diagonal matrix 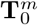 associated with the probability of absence of coancestry change within each of *K* = *L* + 1 hidden states; ii) a matrix 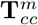 associated with the probability of coancestry change within each state; and iii) a matrix **T**_*cs*_, that does not depend on the marker position, specifying the probability of entering each state after a coancestry change given the state of origin:

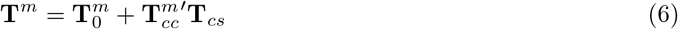

In the nested 1R model, the matrix **T**^**m**^ will have a similar structure but the matrices 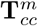 and **T**_*cs*_ in eq. 6 that are defined with respect to states (eq 4) are replaced by matrices 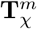 and **T**_*C*_ that are rather defined with respect to layers as we detail below. As a result, **T**^**m**^ is decomposed as:

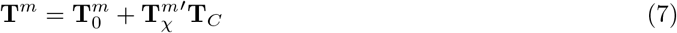

#### 2.2.1 Transition probabilities in nested 1R models

At marker position *m*, the genome can be associated with any state *c* from 1 to *K*. States from 1 to *K −* 1 correspond to HBD segments in layers 1 to *K −* 1 (*K −* 1 being equal to *L*), respectively. HBD segments from state *c* are also non-HBD in layers 1 to *c −* 1. The last state *K* is associated to non-HBD positions in the last layer *L*, and must also be non-HBD in layers 1 to *L −* 1. To estimate the transition probabilities between the *K* different hidden states, we must consider several possible events:

1. the Markov chain remains in the same state *c* without any coancestry change. This requires no coancestry change between the two consecutive markers in all the generations included in both the genealogical paths to the ancestors from layer *c*;
2. the first coancestry change in time along the genealogy occurs within a given layer *c* (i.e., no coancestry change occurs before this layer, in previous generations). We must then account for both the probability of first coancestry change occurring in *c* and the conditional transition probabilities to the other states.

In this model, a coancestry change in layer *c* can be viewed as at least one recombination occurring in that specific layer of ancestors.

#### 2.2.2 Absence of coancestry change from layers 1 to *c*

In the absence of coancestry change between the two consecutive markers, a HBD segment from a given layer *c* is simply extended. The same holds for non-HBD regions in layer *L* (i.e., for the state *K*). The probability of no coancestry change between markers *m* and *m* + 1 from layers 1 to *c* is equal to 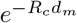, as for a 1R model with rate *R*_*c*_ (eqn 1). These transitions can be summarized for all states as a diagonal matrix 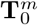:

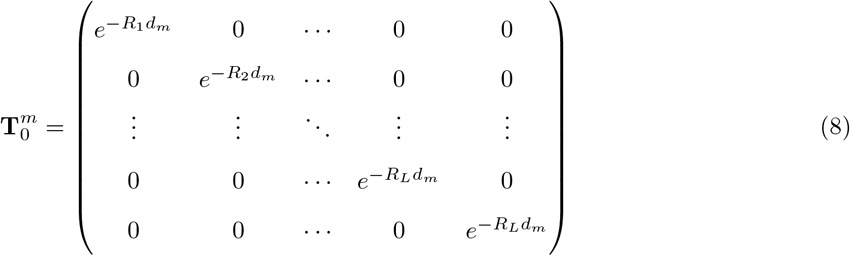

Note that the probabilities for the last two states (*K −* 1 and *K*) are the same as they both belong to the last layer *L*.

#### 2.2.3 Probability of first coancestry change occurring within a given layer *c*

From equation 8, the probability of at least one coancestry change occurring between two consecutive markers *m* and *m* + 1 in the past generations covered by layers 1 to *c* is 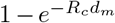. This is in agreement with eqn. 1 for a 1R model with rate *R*_*c*_. The coancestry change may have occurred in any layers *c*′ (1 *≤ c*′ *≤ c*) but we are interested in the first coancestry change event since it implies the start of a new HBD or non-HBD segments in that layer (and thus also affects the status in subsequent layers).

The probability 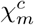 of a first coancestry change occurring within a specific layer *c* is equal to the probability of no coancestry change in earlier layers 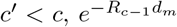, multiplied by the probability of a coancestry change between layers *c −* 1 and *c* which is equal to 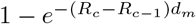:

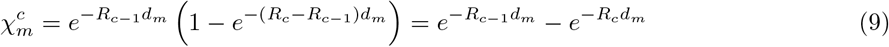

Note that 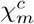 is also the probability of no coancestry change from layer 1 to *c−*1 minus the probability of no coancestry change from layer 1 to *c*. For notational convenience we set *R*_0_ = 0 (i.e., the probability of no coancestry change before the first layer is equal to 1). We can further show that the sum of probabilities of first coancestry changes within each layer from 1 to *c* is equal to 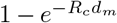 as expected:

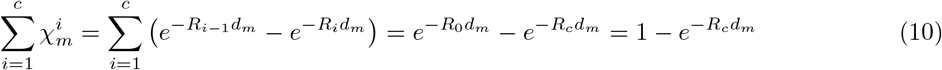

These probabilities can also be combined in a matrix 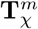, with *K* columns (for states) and *L* rows (for layers). The element 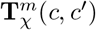 represents the probability of first coancestry change within each layer *c* for a genomic position in an hidden state *c*′ (which is an HBD segment if *c*′ *≤ L* and a non-HBD segment if *c*^′^ = *K*):

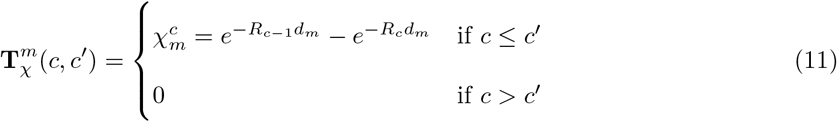

The two last columns of 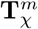 both correspond to probabilities of first coancestry changes for genomic positions in states from the last layer, respectively HBD and non-HBD, and are thus identical. When 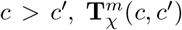 is 0 because HBD segments from layer *c*′ are excluded from more ancestral layers in the modelling (as illustrated in Figure 1). In other words, for a HBD segment in layer *c*′, coancestry changes can only occur from layers 1 to *c*′ because historical crossovers in more remote generations cannot interrupt an HBD tract. Thus, 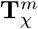 can be represented as:

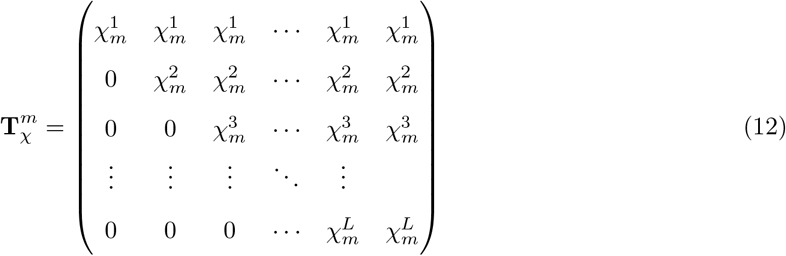

As indicated in Eq. 10, elements from the column *c*′ of 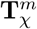 sum to 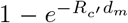 for *c*′ *≤ L*. Each column corresponds to the marginal probability of a coancestry change when the marker *m* is in state *c*′.

#### 2.2.4 Conditional transition probabilities after a coancestry change in layer *c*

If a (first) coancestry change occurs along the genealogy within a given layer *c*, the next position is either i) HBD of class *c* with probability *ρ*_*c*_; or ii) non-HBD with probability 1 *− ρ*_*c*_ (Figure 2). These latter non-HBD regions from layer *c* are also mixture of HBD and non-HBD segments of layer *c* + 1 with probabilities *ρ*_*c*+1_ and 1 *− ρ*_*c*+1_, respectively. The conditional transition probabilities towards the different HBD states *c*′ *> c* and the final non-HBD state from the last layer (i.e., state *K* of the HMM) can then be recursively obtained by following the decision tree represented in Figure 2 (see also Figure 3 for an example of a transition towards the fourth HBD state after a coancestry change in layer *c* = 2). Note that conditional transition probabilities to states *c*′ *< c* that are not a child of the corresponding node in the decision tree (Figure 2) are null. Thus, the conditional transition probabilities **T**_*C*_ (*c, c*′) to reach state *c*′ after a coancestry change occurring in layer *c* are:

**Figure 2.**
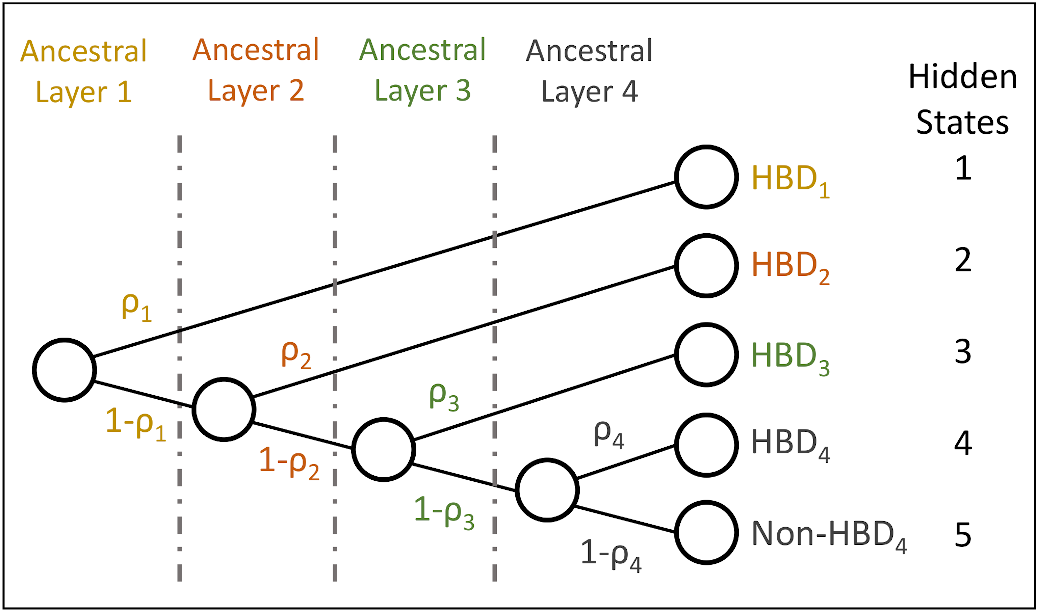
Representation of the transition probabilities in a Nested 1R model with *L* = 4 HBD states and one non-HBD states as a decision tree. In this representation the *K* = 5 states are the leaves of the tree. The tree allows one to estimate probabilities and conditional probabilities to reach a leaf.

**Figure 3.**
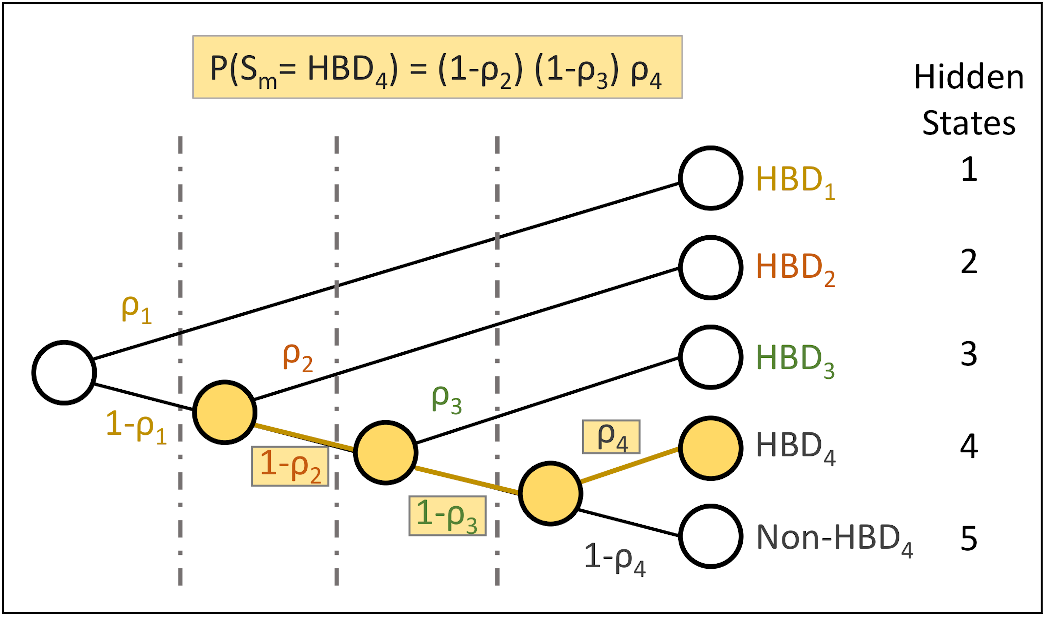
Illustration of conditional transition probabilities after a coancestry change. The illustration shows the conditional transition probability to reach the fourth HBD state after a coancestry change occurring within the second layer.

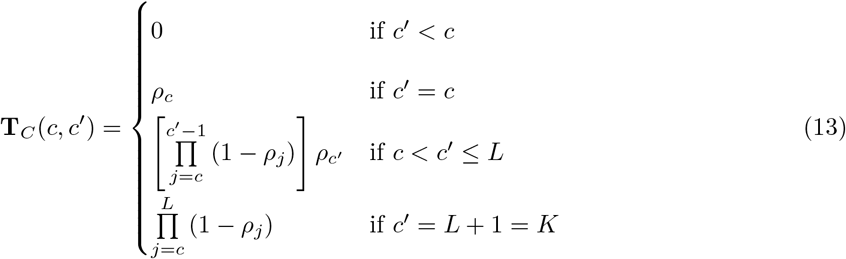

These conditional transition probabilities can be represented as a matrix **T**_*C*_(*c, c*′), independent of the marker position *m*, with *L* rows corresponding to layers, and *K* columns corresponding to the hidden states (*K −* 1 HBD states and one non-HBD state):

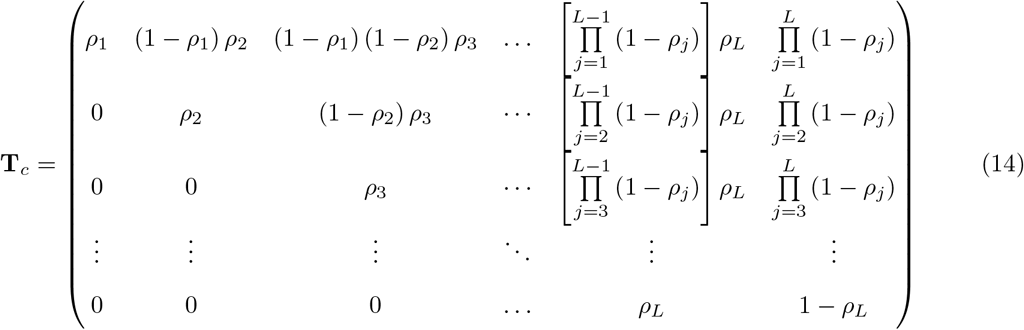

#### 2.2.5 Initial state probabilities

The first row of **T**_*C*_ (eqns. 14 and 7) also corresponds to the vector of initial states probabilities {*δ*_*c*_} _1,…,*K*_ (i.e., *δ*_*c*_ representing the probability to start the chain in the hidden state *c*) which can obtained from the full decision tree (e.g., Figure 2). We have:

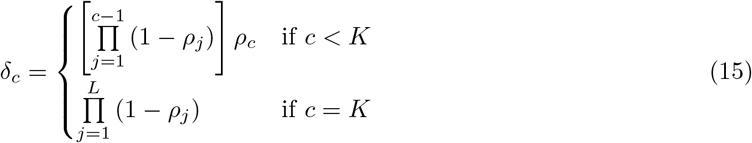

We show in Appendix that the product ***δ*T**^*m*^ = ***δ***, i.e., the Markov chain is stationary and the initial state distribution corresponds to the stationary distribution. These desired properties are also true for the 1R model, but not in our previous MixKR model. Note also that, if *L* = 1, the Nested 1R model reduces to the 1R model.

#### 2.2.6 Parameter estimation

HMM are fully defined by their number of states, their initial state probabilities, their transition probabilities and their emission probabilities; elements that were described in previous sections. However, some of these probabilities depend on the model parameters (*ρ*_*c*_, and *R*_*c*_ for KR models) that need to be estimated. In HMM, this parameter estimation can be performed by numerical maximization of the likelihood with respect to these parameters (Zucchini and MacDonald, 2009). The likelihood of the HMM for a set of parameters is obtained by applying the forward algorithm (Rabiner, 1989). The numerical maximization can be achieved with numerical optimization methods implemented in R packages such as the optim function from the R package stats (R Core Team, 2013). These methods require only a function that returns the likelihood of the HMM for a given set of parameters. In the RZooRoH package, the L-BFGS-B optimizer was selected for that purpose. Following our previous work (Bertrand *et al*., 2019), we transformed the original parameters into new unconstrained parameters as advised by Zucchini and MacDonald (2009). For the rates *R*_*c*_, the following transformation was applied:

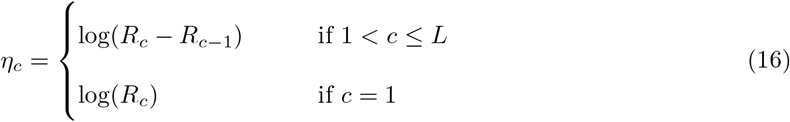

Back-transformation on the original scale ensures that rates are always positive and ordered (higher rates for older ancestral layers). Indeed, as the exponential function is always positive we obtain increasing rates:

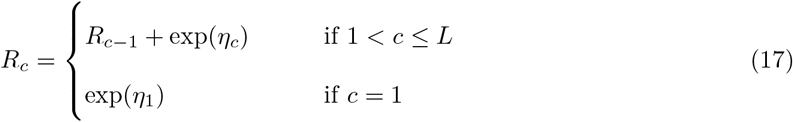

For the the mixing coefficients *ρ*_*c*_, we applied the same transformation as for the MixKR model (Bertrand *et al*., 2019), which here results in a logit transformation (i.e., 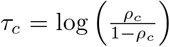 for all *c* ≤ *L*) and guarantees that all the estimated *ρ*_*c*_ are comprised between 0 and 1 (since 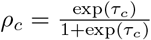).

Note that it may also be possible to rely on numerical optimization with constraints instead of transforming the parameters but this is generally not recommended as the constraints might slow down convergence (Zucchini and MacDonald, 2009). We previously tested different optimization functions and obtained the best results with the parameter transformation and optim L-BFGS-B algorithm.

The N1R model is now implemented as the default model in the RZooRoH package (from version 0.3.1).

#### 2.2.7 Estimation of the inbreeding coefficient

Following Leutenegger *et al*. (2003), the stationary distribution of the state probabilities ***δ*** (eq. 15) can be used to estimate the inbreeding coefficient. We must first define a reference population by deciding which HBD classes are considered as truly autozygous. We could for instance consider that only layers with a rate *R*_*c*_ smaller than a threshold *T* contribute to autozygosity, and that ancestors in layers with *R*_*c*_ *> T* are unrelated (see for instance in Solé et al., 2017). The inbreeding coefficient with respect to that base population, set approximately 0.5*×*T generations in the past (Druet and Gautier, 2017), is:

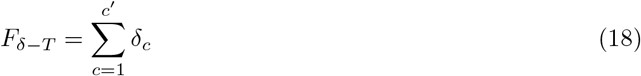

where *c*′ is the most ancient layer with rate *R*_*c*′_ ≤ T. The inbreeding coefficient obtained with all layers is:

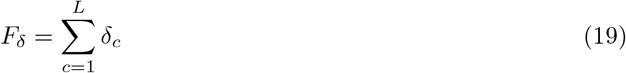

In addition, as opposed to our previous MixKR models, the nested 1R model allows to estimate inbreeding coefficients within each layer. Indeed, the equilibrium probability *ρ*_*c*_ may directly be interpreted as the inbreeding coefficient of the progeny of individuals from the most recent generation of the layer *c* when individuals from the oldest generation of layer *c* are assumed unrelated. This coefficient may also be interpreted as the inbreeding accumulated within the time period covered by layer *c* and may thus be related to the effective population size over this same period. Contrary to the proportion of the genome associated to a specific HBD class, this measure is independent of inbreeding generated in more recent generations.

Metrics defined for the previous MixKR model (Druet and Gautier, 2017) and associated to the realized inbreeding have also their counterpart in the new Nested 1R model. First, the realized inbreeding 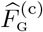 associated with each HBD class *c* (*c ∈* (1, *K −* 1)) can be defined as the proportion of the genome belonging to the class *c* and is estimated as the average of the corresponding local state probabilities over all the *M* loci:

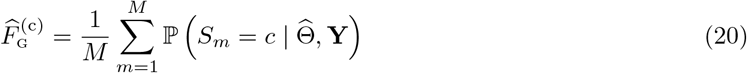

where 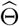 and **Y** represent respectively the estimated parameters of the model and the data.

Next, the genome-wide estimate of the realized individual inbreeding 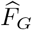 is simply the average over the genome of the local estimates obtained for the *M* markers:

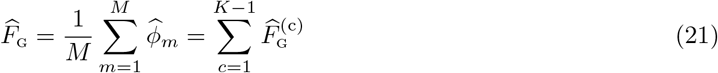

The realized inbreeding coefficients can also be estimated relative to different base populations by considering HBD classes with a rate *R*_*c*_ *≤ T* as in Solé *et al*. (2017).

### 2.3 Evaluation based on simulated data sets

#### 2.3.1 Simulations under the 1R model

To simulate data sets under the 1R model, we used the same approach as in our first study (Druet and Gautier, 2017). Briefly, we simulated individual genomes consisting of 25 chromosomes of 100 cM. Each individual genome is modeled as a mosaic of HBD and non-HBD segments modelled under the 1R model (Equation 1), where *ρ* represents the proportion of HBD segments (equivalent to *F*, the inbreeding coefficient). The length of HBD and non-HBD segments was exponentially distributed with rate *R*. The tested values for *ρ* were equal to 0.02, 0.05, 0.10, 0.20, 0.30 and 0.40, and those for *R* equal to 4, 8, 16, 32 and 64. Genotypes were simulated for 25,000 bi-allelic SNPs (10 per cM) using emission probabilities (Equations 2 and 3). For each set of parameters, we simulated 500 individuals. More details are available in Druet and Gautier (2017).

Individual inbreeding levels were estimated with MixKR and N1R models with 9 HBD classes with rates equal to {2, 4, 8, …, 512}, and using the RZooRoH package (Bertrand *et al*., 2019). The mean absolute error (MAE) for each parameter of interest *α* (*F*_*G*_, *F*_*δ*_, *φ*) was computed to evaluate the models as:

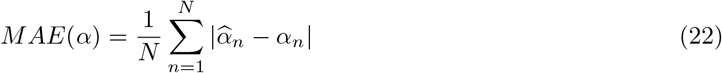

where *N* is the number of simulated individuals, 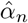 is the estimated parameter value for individual *n* and *α*_*n*_ is the corresponding simulated value.

The partitioning of the autozygosity in different HBD classes was evaluated by assessing whether the autozygosity was concentrated in HBD classes with rates *R*_*c*_ close to the simulated rate *R*. Rates were compared on a log_2_ scale, resulting in a difference of -1, 0, 1 and 2 when *R*_*c*_ is equal to *R* multiplied by respectively 0.5, 1, 2 and 4. The associated MAE was estimated as follows:

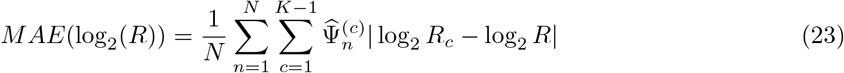

where 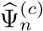 is the contribution of HBD class *c* in individual *n* to its total HBD (computed as 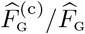 estimated at true HBD positions), and *K −* 1 is the number of HBD classes. This criteria evaluates whether the identified HBD positions are assigned to the simulated HBD class.

#### 2.3.2 Simulations under a discrete-time Wright–Fisher process

To simulate more realistic data relying on population genetic models, Druet and Gautier (2017) previously used the program ARGON (Palamara, 2016) that implements a discrete-time Wright-Fisher process. Here, we used the same simulated data sets. Bottlenecks were simulated to concentrate inbreeding in specific age classes (Druet and Gautier, 2017). Outside these events, *N*_*e*_ was kept large to reduce the noise due to inbreeding coming from other generations. The main simulation scenario is summarized in Supplementary Figure 1. The ancestral population *P*_0_ had a constant haploid effective population size equal to 20,000 (*N*_*e*0_). The time of population split *T*_*s*_ was set equal to 10,000 and the effective population size of the first population (*P*_1_) outside the bottleneck was set to 100,000 (*N*_*e*1_). Bottlenecks were simulated around generations *T*_*b*_ equal to either 16 or 64, and with effective population size (*N*_*eb*_) equal to 20 or 50. A single chromosome of 250 cM length was simulated for 50 diploid individuals, and with a marker density of 100 SNPs per cM. More details about the simulation procedure are available in Druet and Gautier (2017). We further simulated data sets under three additional scenarios to evaluate more complex demographic history. The first two scenarios consisted of two successive bottlenecks (with *N*_*eb*_ being either equal to 20 or 50) either closely related (around generations 16 and 64) or more distant (around generations 16 and 128). The third scenario consisted of a continuous population expansion following a bottleneck simulated around generation 16, the final *N*_*e*_ being ten times larger than the bottleneck one.

Individual inbreeding levels were estimated with MixKR and N1R models with 13 HBD classes with rates equal to {2, 4, 8, …, 8192} as implemented within the RZooRoH package (Bertrand *et al*., 2019).

#### 2.3.3 Application to estimation of inbreeding levels in the European bison

The N1R model was tested and compared to the MixKR model on a set of 183 genotyped European bison with high inbreeding levels (Druet *et al*., 2020). These consisted of respectively 154 and 29 individuals from the Lowland and Lowland-Caucasian lines. Individuals from the first line experienced a stronger bottleneck as they trace back to fewer founders (see Druet *et al*., 2020, for more details). After excluding monomorphic SNPs, those with a call-rate *<*0.95 or deviating from Hardy-Weinberg equilibrium (p *≤* 0.001), each individual was genotyped for 22,602 autosomal SNPs (see Druet *et al*. (2020) for more details). This represents a marker-density below 10 SNPs per cM but we evaluated based on simulations and whole-genome sequence data that it allowed to capture recent autozygosity (Druet *et al*., 2020). Partitioning of inbreeding levels in different HBD classes was first compared with MixKR and N1R models with five HBD classes with rates equal to {4, 8, 16, 32, 64}. In order to assess robustness of results to model specifications, we also applied models with 9 HBD classes (*R*_*c*_ = {4, 8, …, 1024}). Analyses were carried out with the RZooRoH package (Bertrand *et al*., 2019).

## 3 Results

### 3.1 Simulations under the 1R model

We begin by comparing results obtained from analyses of the data simulated under the 1R model with the MixKR and N1R models. We expected our new N1R model to perform better in partitioning inbreeding in different HBD classes most particularly when inbreeding levels are high. This is confirmed in Figure 4 that represents the MAE associated with *R* (eq. 23). With the N1R model, the MAE is higher when there are fewer segments to estimate parameters (small *ρ* and/or small *R*). When inbreeding levels are low (e.g., *ρ <* 0.1 in Figure 4), MAE are similar for both models whereas for large inbreeding levels, MAE starts to increase for the MixKR model whereas it continues to decrease for the N1R model. As a result, the proportions of true HBD positions associated with the class with *R*_*c*_ corresponding to the simulated value is higher with the N1R than with the former MixKR model when values of *ρ* are moderate to high (i.e., *ρ >* 0.1). In other words, a higher proportion of true autozygosity is correctly associated to its underlying HBD class of origin with the N1R model (see Supplementary Table 1). More precisely, these proportions range from 33% up to 84% and actually increased with *ρ* and *R* (i.e., the number of HBD segments available for parameter estimation). Note that in case of mis-classification, the HBD segments are most often assigned to neighboring classes as illustrated for instance in Supplementary Figure 2. As for the MAE, these proportions decrease for high values of *ρ* with the MixKR model whereas an opposite trend is observed for the N1R model, resulting in high differences. When *ρ* is high, the MixKR model tends to assign autozygosity to classes with smaller *R*_*c*_ rates, as we observed in real data sets. This is illustrated for four scenarios with *R* = 8 in Supplementary Figure 2. Similar patterns are obtained in simulations with two distributions of HBD segments (Supplementary Figure 3), with a shift towards more recent HBD classes when using the MixKR model. Finally for comparison with ROH-based methods, we also identified ROH with likelihood-based approaches (Pemberton *et al*., 2012; Szpiech *et al*., 2017) and then assigned these ROH to different classes equivalent to our HBD-classes (see an example in Supplementary Figure 2). We found that the ROH-class that captured the largest proportion of the genome was not the ROH-class including segments of length L=100/2G (the expected length of ROH segments). With higher simulated inbreeding levels, the distribution in different ROH-classes changed and a larger proportion of the genome was found associated with even longer ROH. The inferred distribution of ROH segments was thus sensitive to the overall inbreeding levels as for our previous MixKR model.

**Table 1.**
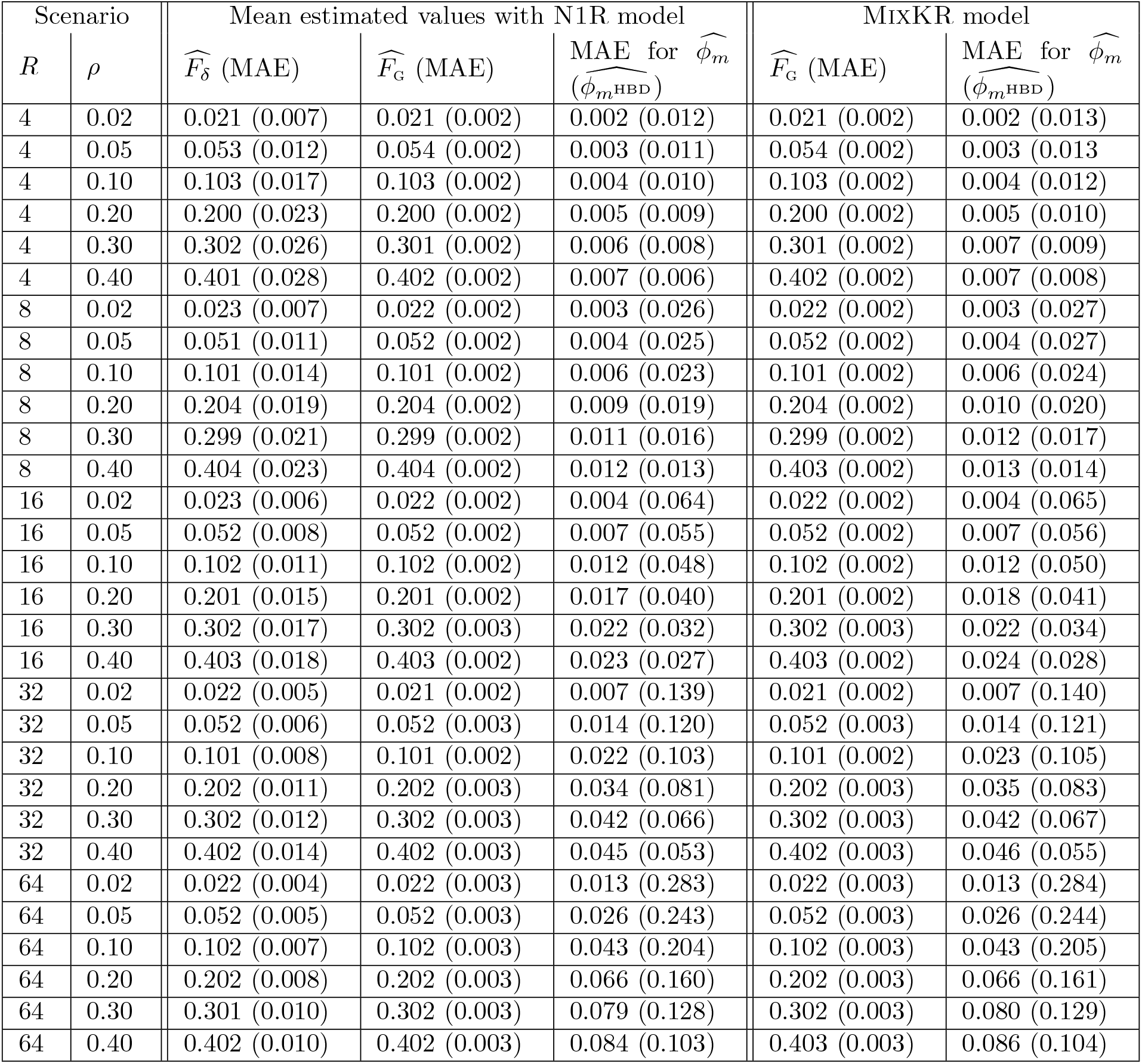
Performance of the two models on data simulated under the 1R model. The simulated genome consisted of 25 chromosomes of 100 cM with a marker density of 10 SNPs per cM. Genotyping data for 500 individuals were simulated under the 1R model for each of 30 different scenarios defined by the simulated *R* and *ρ* values reported in the first two columns. The table reports the mean estimated values and the Mean Absolute Errors (MAE) for the mixing proportions *ρ* estimated as 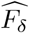 and the individual realized inbreeding levels 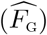. The table gives also the MAE for the estimated local inbreeding (*φ*_*m*_) either for all the genotypes 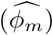 or for genotypes at HBD positions 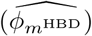. These values are reported for both models, with the exception of 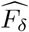.

| Scenario |  | Mean estimated values with N1R model |  |  | MixKR model |  |
| --- | --- | --- | --- | --- | --- | --- |
| $R$ | $\rho$ | $\widehat{F}_\delta$ (MAE) | $\widehat{F}_G$ (MAE) | MAE for $\widehat{\phi}_m$<br>($\widehat{\phi}_{m^{\text{HBD}}}$) | $\widehat{F}_G$ (MAE) | MAE for $\widehat{\phi}_m$<br>($\widehat{\phi}_{m^{\text{HBD}}}$) |
| 4 | 0.02 | 0.021 (0.007) | 0.021 (0.002) | 0.002 (0.012) | 0.021 (0.002) | 0.002 (0.013) |
| 4 | 0.05 | 0.053 (0.012) | 0.054 (0.002) | 0.003 (0.011) | 0.054 (0.002) | 0.003 (0.013) |
| 4 | 0.10 | 0.103 (0.017) | 0.103 (0.002) | 0.004 (0.010) | 0.103 (0.002) | 0.004 (0.012) |
| 4 | 0.20 | 0.200 (0.023) | 0.200 (0.002) | 0.005 (0.009) | 0.200 (0.002) | 0.005 (0.010) |
| 4 | 0.30 | 0.302 (0.026) | 0.301 (0.002) | 0.006 (0.008) | 0.301 (0.002) | 0.007 (0.009) |
| 4 | 0.40 | 0.401 (0.028) | 0.402 (0.002) | 0.007 (0.006) | 0.402 (0.002) | 0.007 (0.008) |
| 8 | 0.02 | 0.023 (0.007) | 0.022 (0.002) | 0.003 (0.026) | 0.022 (0.002) | 0.003 (0.027) |
| 8 | 0.05 | 0.051 (0.011) | 0.052 (0.002) | 0.004 (0.025) | 0.052 (0.002) | 0.004 (0.027) |
| 8 | 0.10 | 0.101 (0.014) | 0.101 (0.002) | 0.006 (0.023) | 0.101 (0.002) | 0.006 (0.024) |
| 8 | 0.20 | 0.204 (0.019) | 0.204 (0.002) | 0.009 (0.019) | 0.204 (0.002) | 0.010 (0.020) |
| 8 | 0.30 | 0.299 (0.021) | 0.299 (0.002) | 0.011 (0.016) | 0.299 (0.002) | 0.012 (0.017) |
| 8 | 0.40 | 0.404 (0.023) | 0.404 (0.002) | 0.012 (0.013) | 0.403 (0.002) | 0.013 (0.014) |
| 16 | 0.02 | 0.023 (0.006) | 0.022 (0.002) | 0.004 (0.064) | 0.022 (0.002) | 0.004 (0.065) |
| 16 | 0.05 | 0.052 (0.008) | 0.052 (0.002) | 0.007 (0.055) | 0.052 (0.002) | 0.007 (0.056) |
| 16 | 0.10 | 0.102 (0.011) | 0.102 (0.002) | 0.012 (0.048) | 0.102 (0.002) | 0.012 (0.050) |
| 16 | 0.20 | 0.201 (0.015) | 0.201 (0.002) | 0.017 (0.040) | 0.201 (0.002) | 0.018 (0.041) |
| 16 | 0.30 | 0.302 (0.017) | 0.302 (0.003) | 0.022 (0.032) | 0.302 (0.003) | 0.022 (0.034) |
| 16 | 0.40 | 0.403 (0.018) | 0.403 (0.002) | 0.023 (0.027) | 0.403 (0.002) | 0.024 (0.028) |
| 32 | 0.02 | 0.022 (0.005) | 0.021 (0.002) | 0.007 (0.139) | 0.021 (0.002) | 0.007 (0.140) |
| 32 | 0.05 | 0.052 (0.006) | 0.052 (0.003) | 0.014 (0.120) | 0.052 (0.003) | 0.014 (0.121) |
| 32 | 0.10 | 0.101 (0.008) | 0.101 (0.002) | 0.022 (0.103) | 0.101 (0.002) | 0.023 (0.105) |
| 32 | 0.20 | 0.202 (0.011) | 0.202 (0.003) | 0.034 (0.081) | 0.202 (0.003) | 0.035 (0.083) |
| 32 | 0.30 | 0.302 (0.012) | 0.302 (0.003) | 0.042 (0.066) | 0.302 (0.003) | 0.042 (0.067) |
| 32 | 0.40 | 0.402 (0.014) | 0.402 (0.003) | 0.045 (0.053) | 0.402 (0.003) | 0.046 (0.055) |
| 64 | 0.02 | 0.022 (0.004) | 0.022 (0.003) | 0.013 (0.283) | 0.022 (0.003) | 0.013 (0.284) |
| 64 | 0.05 | 0.052 (0.005) | 0.052 (0.003) | 0.026 (0.243) | 0.052 (0.003) | 0.026 (0.244) |
| 64 | 0.10 | 0.102 (0.007) | 0.102 (0.003) | 0.043 (0.204) | 0.102 (0.003) | 0.043 (0.205) |
| 64 | 0.20 | 0.202 (0.008) | 0.202 (0.003) | 0.066 (0.160) | 0.202 (0.003) | 0.066 (0.161) |
| 64 | 0.30 | 0.301 (0.010) | 0.302 (0.003) | 0.079 (0.128) | 0.302 (0.003) | 0.080 (0.129) |
| 64 | 0.40 | 0.402 (0.010) | 0.402 (0.003) | 0.084 (0.103) | 0.403 (0.003) | 0.086 (0.104) |

**Figure 4.**
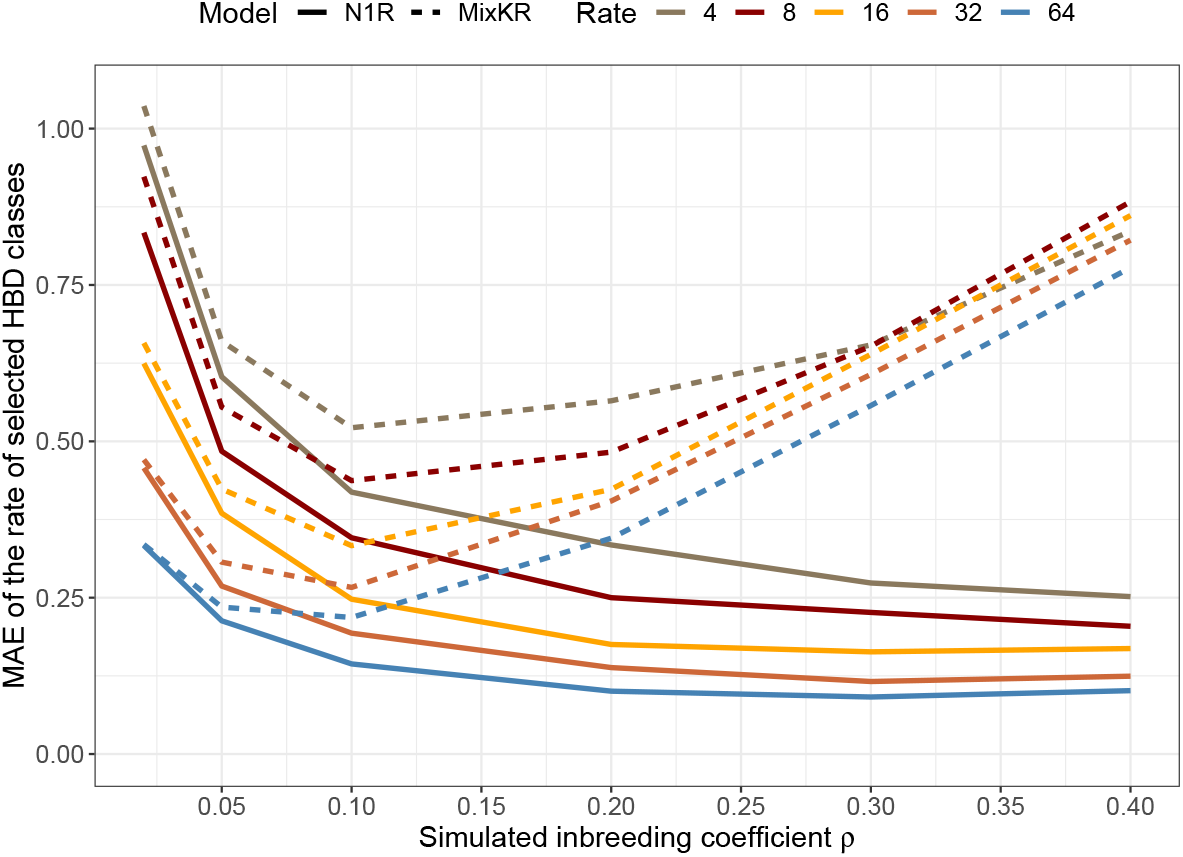
Concordance between simulated rates and partitioning in HBD classes on data sets simulated under the 1R model. The accuracy of partitioning is evaluated as the Mean Absolute Error between the log_2_ of the simulated rate and the log_2_ of the assigned HBD classes. This is equivalent to measuring the deviation from the simulated parameter in term of absolute value of log_2_ of the ratio between rates of simulated and estimated HBD classes. The comparisons are performed for different values of *R* and *ρ*.

In terms of estimation of realized inbreeding (*F*_*G*_) and estimation of local HBD probabilities (*φ*_*m*_), both the MixKR and N1R models showed very similar performances (Table 1). Hence, although these two models differ in their partitioning of inbreeding in different age classes, they remain equally accurate for the estimation of inbreeding levels. Finally, the inbreeding coefficient *F*_*δ*_ corresponding to the sum of initial state probabilities (the stationary distributions) and closely related to *ρ* displayed a low MAE, close to the values obtained with a 1*R* model in our previous study (Druet and Gautier, 2017). With the N1R model, the inbreeding coefficient *F*_*δ*_ represents an unbiased estimate of the simulated *ρ* (Supplementary Figure 4), as opposed to the sum of initial state probabilities of HBD classes from the MixKR models that were clearly not a proper estimator of *ρ*.

### 3.2 Simulations under Wright-Fisher process

Analyses realized on data sets simulated under a more realistic model confirm our first observations. For high inbreeding levels, the MixKR model captures a large fraction of the autozygosity generated by the bottleneck (when *N*_*e*_ drops to 20) into the more recent HBD class neighboring the class representative of the bottleneck period (e.g., class with *R*_*c*_ = 64 for a bottleneck pertaining to the class with *R*_*c*_ = 128, i.e., occurring 63 to 66 generations ago - Figure 5). This neighbouring class captures almost the same or even a larger fraction of autozygosity that the HBD class associated with the bottleneck. The pattern is less pronounced for milder bottleneck (*N*_*e*_ =50 in Figure 5). With the N1R model, the class *R*_*c*_ = 128 representative of the bottleneck period captures the majority of the HBD segments in both cases. Similar results were obtained for more recent bottlenecks (Supplementary Figure 5).

**Figure 5.**
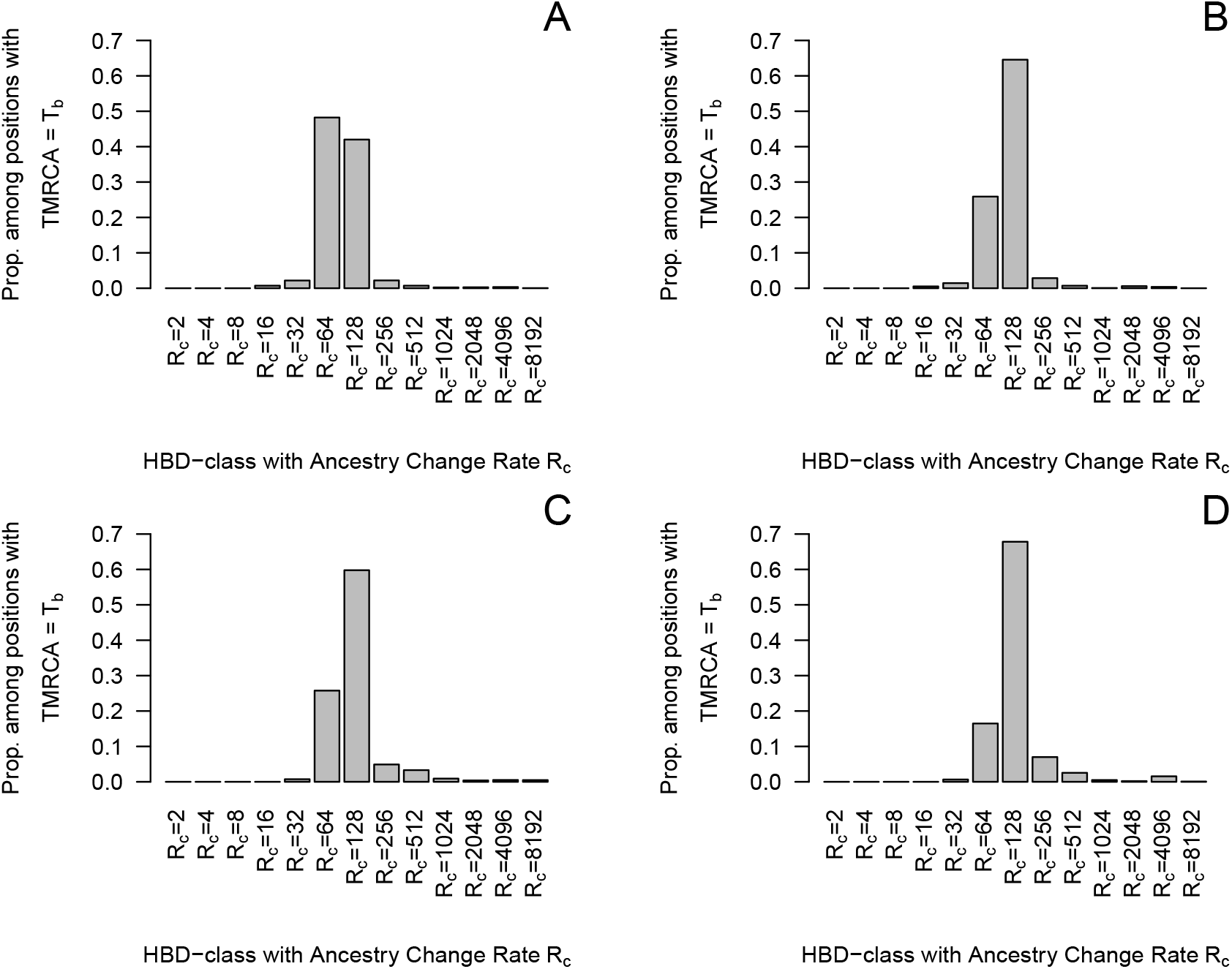
Partitioning of HBD segments related to the bottleneck in different HBD classes. Data were simulated with a Wright-Fisher process, with a bottleneck in generations 63 to 66 expected to be associated with the HBD class with *R*_*c*_ = 128. The simulated *N*_*e*_ during the bottleneck is equal to 20 (A & B) or 50 (C & D). The partitioning is realized with the MixKR (A & C) or N1R (B & D) models.

The global partitioning of the genome in HBD-classes presents similar patterns (Supplementary Figure 6). As the proportion of inbreeding in the HBD class associated with the bottleneck is always higher with the N1R model, the MAE associated with the rate of the selected HBD classes is lower than with the MixKR model (more so when the bottleneck was strong). With the N1R model, the MAE values are respectively equal to 0.546, 0.786, 0.386 and 0.426 for the four different scenarios (*{N*_*eb*_ = 20, *T*_*b*_ = 16},*{N*_*eb*_ = 50, *T*_*b*_ = 16},*{N*_*eb*_ = 20, *T*_*b*_ = 64},*{N*_*eb*_ = 50, *T*_*b*_ = 64}), compared to 0.763, 0.793, 0.601 and 0.491 for the same scenarios with the MixKR model.

**Figure 6.**
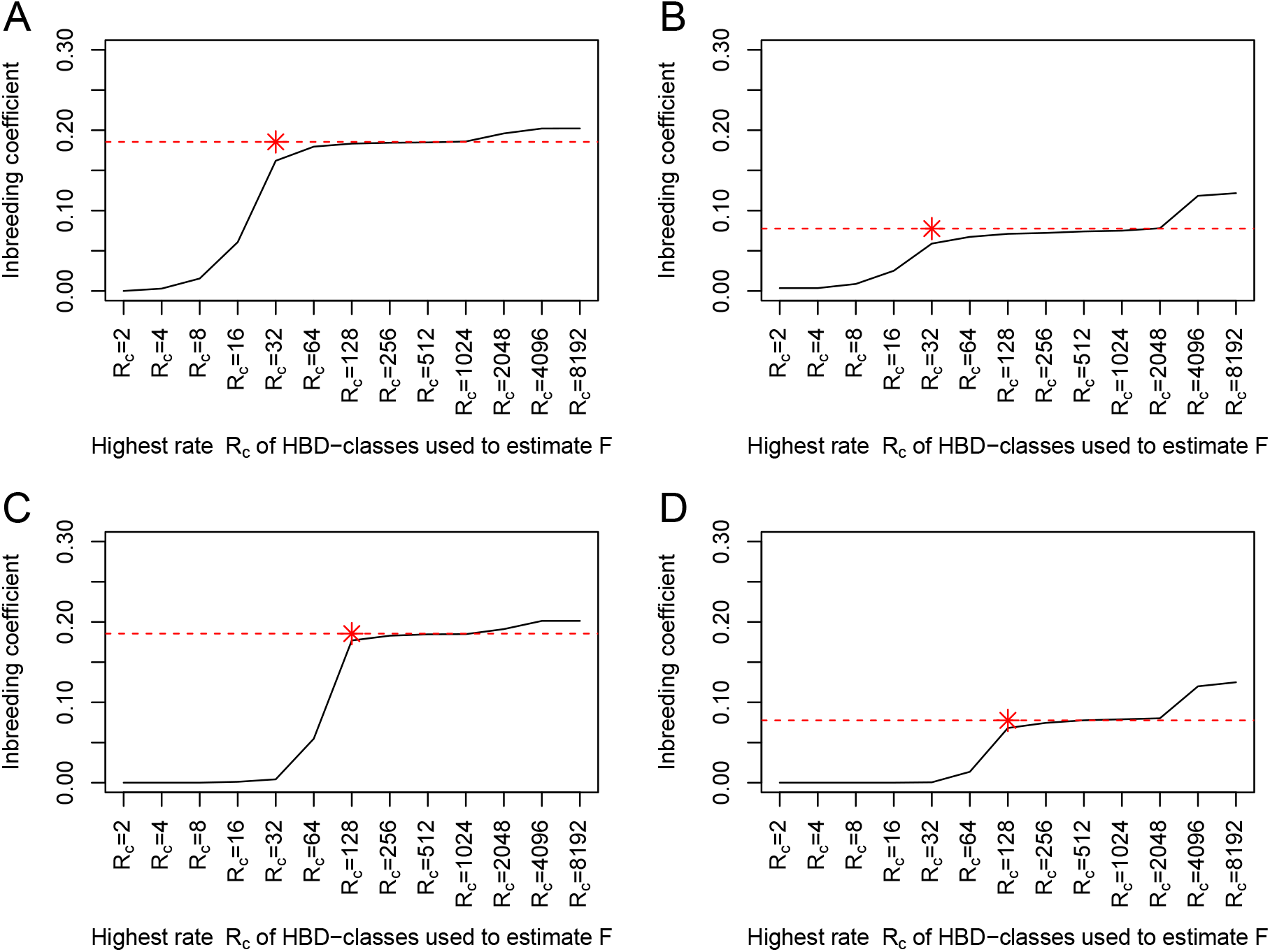
Inbreeding coefficients estimated as the equilibrium HBD distribution and for different base generations. The inbreeding coefficients *F*_*δ−T*_ are estimated from the equilibrium distributions from the different HBD classes and are obtained from their mixing coefficients *ρ*_*c*_ (see eq. 18)). Only HBD-classes with a rate *R*_*c*_ *≤* a threshold T are used to estimate *F*_*δ−T*_. This allows to set the reference population approximately 0.5 *× T* generations in the past. Data were simulated with a Wright-Fisher process, with a bottleneck. The time of the bottleneck (*T*_*b*_) and the effective population size during the bottleneck *N*_*eb*_ are A) *T*_*b*_ = 16 and *N*_*eb*_ = 20, B) *T*_*b*_ = 16 and *N*_*eb*_ = 50, C) *T*_*b*_ = 64 and *N*_*eb*_ = 20, D) *T*_*b*_ = 64 and *N*_*eb*_ = 50. The red star indicates the HBD-class associated to the bottleneck and the expected inbreeding levels generated during the bottleneck.

As for the simulations under the 1R model, the differences between models are mainly in the partitioning of autozygosity in HBD classes. For instance, the average local HBD probabilities for positions associated with ancestors present in different past generations are almost identical (Supplementary Figure 7). These local HBD probabilities indicate that HBD positions associated with ancestors up to 80 generations ago are identified with high confidence, and that for more remote ancestors (shorter segments), it is more difficult to identify unambiguously HBD positions. We also confirm in Figure 6 that mixing coefficients of the new model are interpretable and can be used to estimate the inbreeding coefficient *F*_*δ*_. More precisely, we estimated *F*_*δ−T*_ by adding sequentially each HBD-class in the estimation. We estimated the expected inbreeding accumulated during the bottleneck as 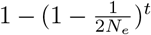, where *N*_*e*_ is the diploid effective population size (here, *N*_*e*_ = 20 or *N*_*e*_ = 50) and *t* = 4 is the number of generations of the bottleneck. We see that most of the inbreeding is captured by the HBD-class corresponding to the bottleneck and its close neighbours. As a result, *F*_*δ−T*_ remains relatively constant for generations before and after the bottleneck and increases sharply during the bottleneck. In addition, the estimated inbreeding levels match the expected values. We also observe inbreeding related to much more distant ancestors, accumulated over many more generations that was captured by the most remote HBD classes (e.g., *R*_*c*_ *≥* 2048). Note that the ability of both models to capture ancient bottlenecks obviously depends on the marker density as we previously showed (Druet and Gautier, 2017). For instance, with a marker density of 10 SNPs per cM, a bottleneck occurring 16 generations ago would be fully captured whereas HBD-segments from a later bottleneck (G=64) would only be partially captured.

**Figure 7.**
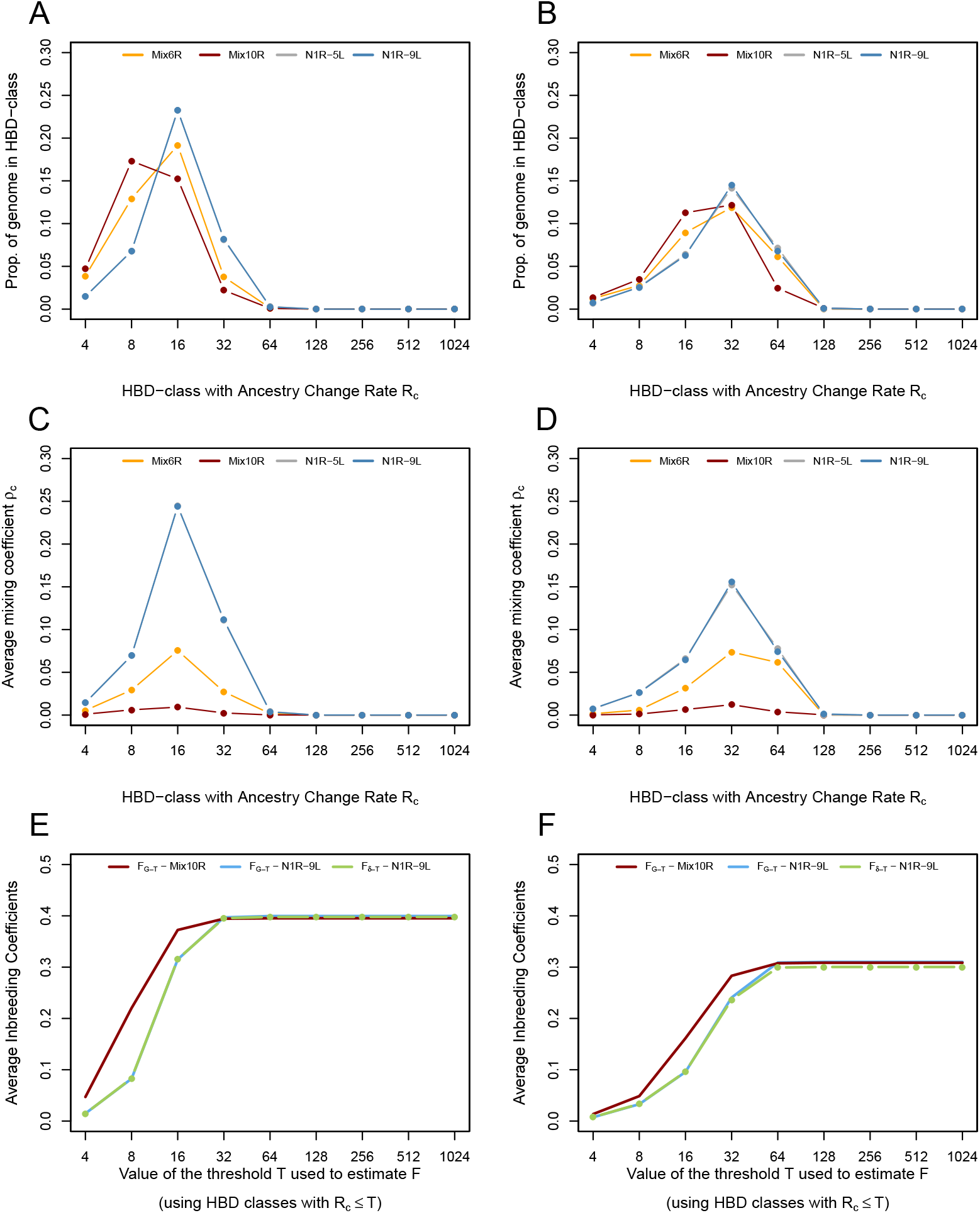
Estimation of inbreeding levels in the European bison. Inbreeding levels are estimated in 154 Lowland individuals (panels A-C-E) and 29 Lowland-Caucasian individuals (panels B-D-F). Estimation was performed with the MixKR and N1R models with 5 HBD-classes (Mix6R and N1R-5L) or with 9 HBD-classes (Mix10R and N1R-9L). A) and B) Proportion of the genome associated with different HBD-classes averaged over all individuals from a population (see eq. 20). C) and D) Estimated mixing coefficients *ρ*_*c*_ for each HBD class, averaged over all individuals. E) and F) Average estimated inbreeding levels. *F*_*G−T*_ is estimated as a the sum of contributions from the HBD classes with a rate *R*_*c*_ *≤* a threshold *T*, and *F*_*δ−T*_ as the sum of equilibrium distributions of the same HBD classes (see eq. 18). This allows to set the reference population approximately 0.5 *× T* generations in the past. A cumulative curve is obtained by changing the values of *T*.

When autozygosity results from more complex demographic histories such as multiple bottlenecks or gradual population expansion after a bottleneck, it becomes more difficult to determine precisely which generations contribute to autozygosity and to classify HBD positions as illustrated in Supplementary Figures 8 to 13. For instance, in the presence of two relatively distant bottlenecks (in number of generations), the inferred contributions of the different HBD classes looks bimodal (Supplementary Figures 8 and 9), with the modes corresponding to the two classes representative of the bottleneck. However, the shortest segments of the recent bottleneck might sometimes be assigned to the older classes and vice versa. When the two bottleneck are closer, we no longer observe clear bimodality of the estimated distribution of HBD contributions, and the highest contribution is associated to a class located between the two classes representative of the bottleneck (Supplementary Figures 10 and 11). We previously and more comprehensively studied the properties of the MixKR model when several classes contributed to autozygosity and observed similar trends, with difficulties to disentangle contributions from close HBD classes (Druet and Gautier, 2017). As expected, in the scenario with population expansion following a bottleneck, the distribution of HBD classes contribution tends to be shifted towards classes representative of more recent times than the bottleneck, i.e., with *R*_*c*_ *<* 32 (Supplementary Figures 12 and 13). Likewise, the shift is more pronounced for smaller *N*_*eb*_ (i.e., for higher bottleneck intensity). This is likely related to the non negligible contribution to inbreeding of generations that immediately follow the bottlenecks (when *N*_*e*_ is still small) (e.g., those directly following the bottleneck which might have a substantial contribution). The classification was indeed better when all the autozygosity was associated to the bottleneck as shown for scenarios with a rapid expansion after the bottleneck (Figure 5 and Supplementary Figure 5). Nevertheless, both models still indicate periods of increased contribution to inbreeding, from the start of bottlenecks until the time period when the population has recovered. Interestingly, the estimated contributions from HBD classes to autozygosity tend to be less shifted towards more recent classes with the N1R than with the MixKR model.

### 3.3 Application to real data

Application of the two models on genotype data from two distinct lines of European bison, presenting high inbreeding levels, results in similar observations than applications to simulated data sets: partitioning of inbreeding in HBD-class is shifted towards more recent HBD-classes with the MixKR model compared to the N1R model (Figure 7A-B). Since for simulations the N1R performed better for the partitioning in HBD-classes, and since patterns are similar, the results from the N1R model fit probably better the reality. The shift is more pronounced when more HBD-classes are included in the MixKR model and the non-HBD class has consequently a higher rate *R*_*K*_, and in the Lowland line where the inbreeding levels are higher. Higher shifts for higher inbreeding levels were also observed with simulated data. With the MixKR model, the partitioning in different HBD-classes and the estimated mixing coefficients (Figure 7C-D) change according to the model specifications, whereas the N1R model proves robust to these changes (Figures 7A-D). Note that we also fitted HBD-classes corresponding to HBD segments shorter than the shortest HBD segments than could be captured with the available density. As a result, the contribution of these classes remains null. As for the simulated data sets, the overall inbreeding levels estimated by MixKR and N1R models remain highly similar (Figure 7E-F), the difference being essentially in the partitioning.

Analysis of real data with the N1R model confirms that mixing coefficients can now be interpreted, with levels close to estimated HBD proportions in different classes, contrary to those obtained with the MixKR model (Figure 7C-D). In addition, they can now be used to estimate the inbreeding coefficients, *F*_*δ*_ or *F*_*δ−T*_. These inbreeding coefficients based on the equilibrium distribution and on the number of HBD segments are close to values of the realized inbreeding coefficient, *F*_*G*_ and *F*_*G−T*_, corresponding to the proportion of the genome in HBD classes (Figures 7E-F). The mixing coefficients estimate the proportion of HBD segments within a specific layer and provide an estimation of the inbreeding accumulated in that layer, which depends also on the number of generations included in the layer.

When inbreeding levels are lower, such as in cattle (see for instance in Solé *et al*. (2017)), differences are smaller. This is illustrated in Supplementary Figure 14 on a Holstein data set including 245 individuals genotyped for 30,000 markers (Alemu *et al*., 2021).

## 4 Discussion

We herein proposed an improved model, we called the N1R model, for the characterization of individual genomic inbreeding levels and its partitioning into different HBD-classes. Compared to our previous MixKR model (Druet and Gautier, 2017), the main improvement relied on a new modelling of the transition probabilities which both resulted in better statistical properties in general, but also facilitated the interpretation of the mixing coefficients with initial state probabilities now corresponding to the stationary distributions. Although the estimation of both global and local inbreeding levels were almost identical between the N1R and the MixKR models, the partitioning of inbreeding into different HBD-classes was clearly improved and the N1R model provided more accurate estimation of the relative contribution of each group of ancestors.

Our main objective was indeed to improve this partitioning, in particular for high inbreeding levels since we previously observed that in such cases, the partitioning could be shifted towards more recent HBD classes (Druet *et al*., 2020). This problem was caused in our previous MixKR model by the difference of rates for HBD classes associated with recent ancestors (i.e., capturing large HBD segments) and the non-HBD class that resulted in high differences in their underlying mixing coefficients. More precisely, the non-HBD class had a very high mixing coefficient because it generally represented the main contribution to individual genomes and it was modelled with a large *R*_*c*_. Conversely, mixing coefficients from recent HBD classes (long segments with low rates *R*_*c*_) were very small as these segments were much less numerous than short HBD segments. Therefore, in the Markov chain, the probability to start a new recent HBD segment was extremely low and needed to be supported by long stretches of homozygous genotypes. In these conditions, two consecutive recent HBD segments were systematically modelled as a single long HBD segments because transitions to new recent HBD segments were heavily penalized, explaining the overestimation of segment length and the incorrect HBD partitioning (a shift towards more recent HBD classes). Yet, the strength of this problem was expected to be a function of the frequency of consecutive HBD segments, and was thus only observed in simulated and real data sets with high recent inbreeding levels (Druet *et al*., 2020). We here showed that using the same rates for HBD and non-HBD states by modelling sequentially multiple nested 1R models in our new N1R model allowed to solve this issue. This property is important to better interpret the results by determining which generations of ancestors mostly contributed to autozygosity. Our improved N1R model should also allow better estimation of the number of generations to the common ancestor for an HBD position. Nevertheless, more work is required to quantify how precisely the age of individual HBD segments can be estimated with this or other similar approaches.

The new model is also more robust to the number and specifications (i.e., rates *R*_*c*_) of the fitted classes in the sense that partitioning remains consistent when the rate of the non-HBD classes is modified. With our previous MixKR model, the choice of the rates associated with the non-HBD classes, often directly related to the number of fitted classes, might indeed influence the partitioning in HBD and non-HBD classes because higher rates (smaller segments) resulted in even higher mixing coefficients for the non-HBD class further penalizing the occurrence of two consecutive recent HBD segments (see above). The fact that the N1R model is less sensitive to model specifications is an important aspect because one of the advantages of methods relying on HMM (Leutenegger *et al*., 2003; Vieira *et al*., 2016; Narasimhan *et al*., 2016; Druet and Gautier, 2017) is that fewer parameters need to be defined compared to rule-based ROH approaches, where these definitions might sometimes be arbitrary. In general, there is less need to optimize parameters, HBD probabilities indicate whether the evidence for autozygosity is strong or not. In our model, the number of classes and their range must still be defined but it affects mainly interpretation in terms of age of ancestors. To this respect, the robustness of the N1R model is highly valuable since in the previous MixKR model partitioning could be affected by the definition of the last HBD class.

Our newly developed N1R model allows the definition of new inbreeding coefficients based on the initial state probabilities. These inbreeding coefficients fit closer to the original definition by Leutenegger *et al*. (2003) since under the 1R model, the mixing coefficient can be interpreted as both the frequency of HBD segment and the proportion of the genome that is HBD (i.e., the equilibrium distribution). Yet, this is slightly different from a direct estimation of the realized proportion of the genome in HBD segments (e.g., as obtained from the posterior HBD probability of each marker, see eq. 21), although both estimators are highly correlated. Interestingly, the mixing coefficients also provide direct estimators of the level of inbreeding associated with ancestors present in a specific period of time (corresponding to a layer in our model), independently on what happened in other more recent layers. In an ideal population, this inbreeding would directly be related to the number of generations and to the effective population size in the layer. These aspects must be further investigated and more work is required to understand which generations are captured by a specific layer, or the relationship with the underlying historical *N*_*e*_. Indeed, generations do not map unambiguously to a single layer but are captured by several layers in a probabilistic framework. In practice, the variation of mixing coefficients across layers could be used to monitor whether inbreeding is increasing or not, for instance in a conservation program as suggested by Druet *et al*. (2020).

Comparisons of our previous MixKR and our new N1R models on genotyping data from European bison were in agreement with trends observed on simulated data. The overall inbreeding levels were similar with both models but the partitioning was different, shifted towards more recent HBD classes with the MixKR model. This shift was also more pronounced when inbreeding levels were higher and when the rate of the non-HBD class was higher, matching our predictions (see above). This suggests that the new partitioning is more accurate, strengthening our initial conclusions that the contribution from the most recent generations of ancestors to inbreeding is decreasing and that the restoration plan has been successful to control inbreeding in European bison (Druet *et al*., 2020).

Finally, it is important to note that differences between our new N1R version of the model and the former MixKR one in terms of interpretation only concern the partitioning of inbreeding when inbreeding levels are high. For instance, differences would be minimal in most human populations. Even in cattle presenting moderate inbreeding levels, the impact on the partitioning remained limited.

## 5 Acknowledgements

We thank the three anonymous reviewers for their valuable and constructive comments that helped us improving the original version of the manuscript. This work was supported by the Fonds de la Recherche Scientifique–FNRS under grants J.0134.16 and J.0154.18. Tom Druet is Senior Research Associate from the F.R.S.–FNRS. We used the supercomputing facilities of the “Consortium d’Equipements en Calcul Intensif en Fédération Wallonie-Bruxelles” (CECI), funded by the F.R.S.–FNRS.

## A Appendix

Here we show that in the N1R model, the Markov chain is stationary and the initial state distribution corresponds to the stationary distribution, i.e.:

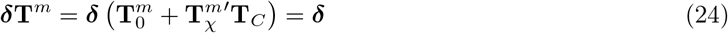

where ***δ*** is a row vector of dimension *K*. Let the (row) vector *ζ* = {*ζ*_**c**_}_**1**,..,**K**_ = ***δ*T**^**m**^. We want to show that 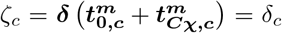 for all c *∈* (1, *K*), where 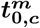 is the *c*th column vector of 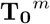 and 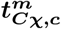 is the *c*th column vector of the matrix 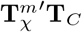:

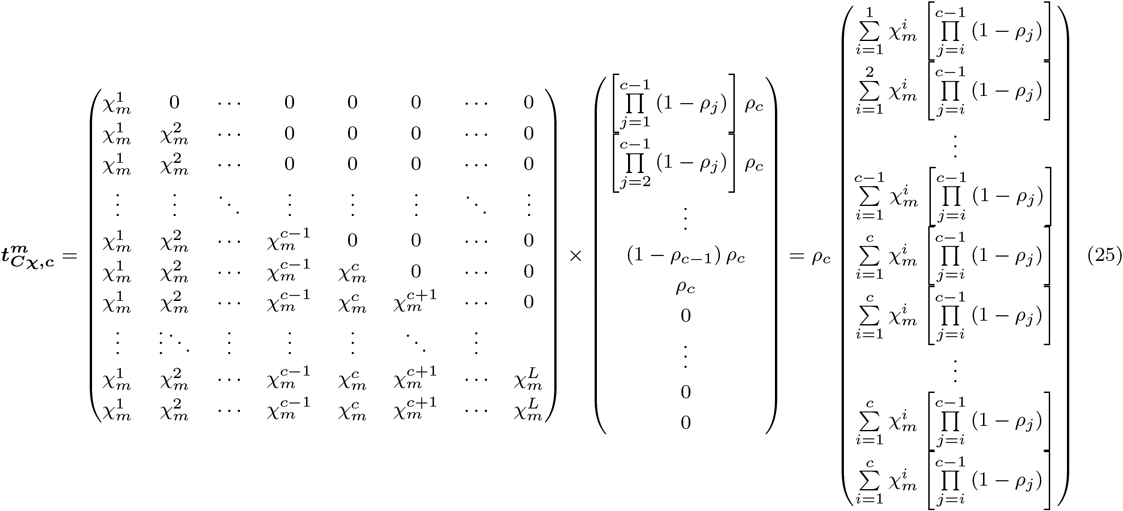

To simplify notations in the above equation, we assume that 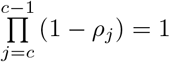. Still to keep notations general, for *c* = *K* we define *ρ*_*K*_ = 1 *− ρ*_*K−*1_. Note also that elements *c*′ ≤ *c* of 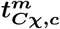 are all identical.

Hence,

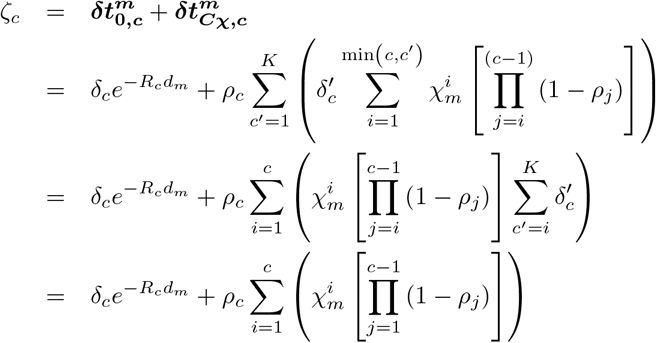

The last equality follows from the nested model properties which consider each layer sequentially (see the main text and Figure 2). Hence, 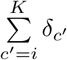 can be interpreted as the probability of starting a layer as old or older than *i* which is also the probability of not having entered any of the successive layers more recent than *i* i.e. 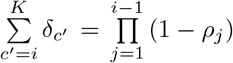. Note also that 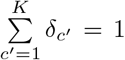. In addition, recalling that 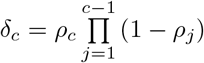 (eq. 15) and 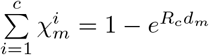 (eq. 10), we obtain:

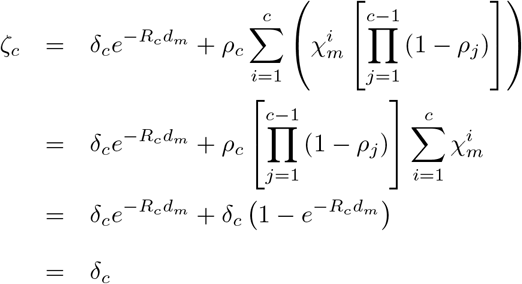

## Notes

### Competing Interest Statement

The authors have declared no competing interest.

