## Supplementary Material for "An hidden Markov model to estimate homozygous-by-descent probabilities associated with nested layers of ancestors"

for

November 15, 2021

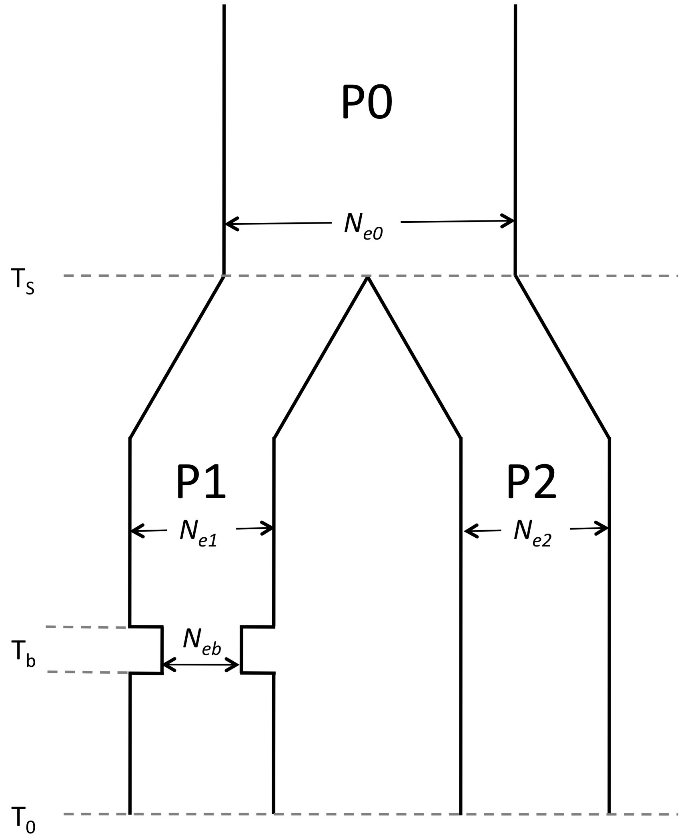


**Supplementary Figure 1. Description of the first simulated scenario under the discrete time Wright-Fisher process (WF1).** We start with an ancestral population $P_0$ with a constant haploid effective population size equal to $N_{e0}$=20,000 that splits in two populations $P_1$ and $P_2$ at generation time $T_s$ in the past with respective population sizes $N_{e1}$=10,000 or 100,000 (according to the scenario) and $N_{e2}$=10,000. During four generations centered around generation $T_b \ll T_s$ in the past, $P_1$ experienced a bottleneck with an (haploid) effective population size equal to $N_{eb}$ and recovered its initial size. Population $P_2$ that always maintains a constant size is actually used to select markers that were also segregating in the ancestral population $P_0$ (markers segregating at MAF $\geqslant$ 0.05 in the present day individuals from both populations $P_1$ and $P_2$ were kept for further analyses). Estimation of inbreeding was performed on 50 diploid individuals from population $P_1$ (in $T_0$) and allele frequencies used by our model were estimated on these individuals.

| Scenario | | Proportion of HBD in the simulated HBD class | |
|---|---|---|---|
| $R$ | $\rho$ | N1R model | MixKR model |
| 4 | 0.02 | 0.328 | 0.296 |
| 4 | 0.05 | 0.492 | 0.433 |
| 4 | 0.10 | 0.602 | 0.486 |
| 4 | 0.20 | 0.670 | 0.412 |
| 4 | 0.30 | 0.728 | 0.324 |
| 4 | 0.40 | 0.748 | 0.155 |
| 8 | 0.02 | 0.421 | 0.375 |
| 8 | 0.05 | 0.581 | 0.504 |
| 8 | 0.10 | 0.667 | 0.583 |
| 8 | 0.20 | 0.761 | 0.524 |
| 8 | 0.30 | 0.778 | 0.369 |
| 8 | 0.40 | 0.812 | 0.184 |
| 16 | 0.02 | 0.486 | 0.455 |
| 16 | 0.05 | 0.632 | 0.590 |
| 16 | 0.10 | 0.732 | 0.641 |
| 16 | 0.20 | 0.807 | 0.554 |
| 16 | 0.30 | 0.824 | 0.359 |
| 16 | 0.40 | 0.830 | 0.167 |
| 32 | 0.02 | 0.525 | 0.507 |
| 32 | 0.05 | 0.670 | 0.630 |
| 32 | 0.10 | 0.736 | 0.663 |
| 32 | 0.20 | 0.799 | 0.544 |
| 32 | 0.30 | 0.835 | 0.369 |
| 32 | 0.40 | 0.840 | 0.180 |
| 64 | 0.02 | 0.504 | 0.500 |
| 64 | 0.05 | 0.603 | 0.584 |
| 64 | 0.10 | 0.690 | 0.625 |
| 64 | 0.20 | 0.762 | 0.552 |
| 64 | 0.30 | 0.798 | 0.385 |
| 64 | 0.40 | 0.811 | 0.202 |

**Supplementary Table 1. Proportion of HBD positions assigned to the simulated HBD classes.** The results are obtained on simulated genotypes for 500 individuals with different scenarios defined by the simulated $R$ and $\rho$ values reported in the first two columns. Estimation with multiple HBD classes models was realized with a MixKR and a N1R model. The reported values are the proportions of true HBD positions correctly assigned to the simulated HBD class, across all individuals.

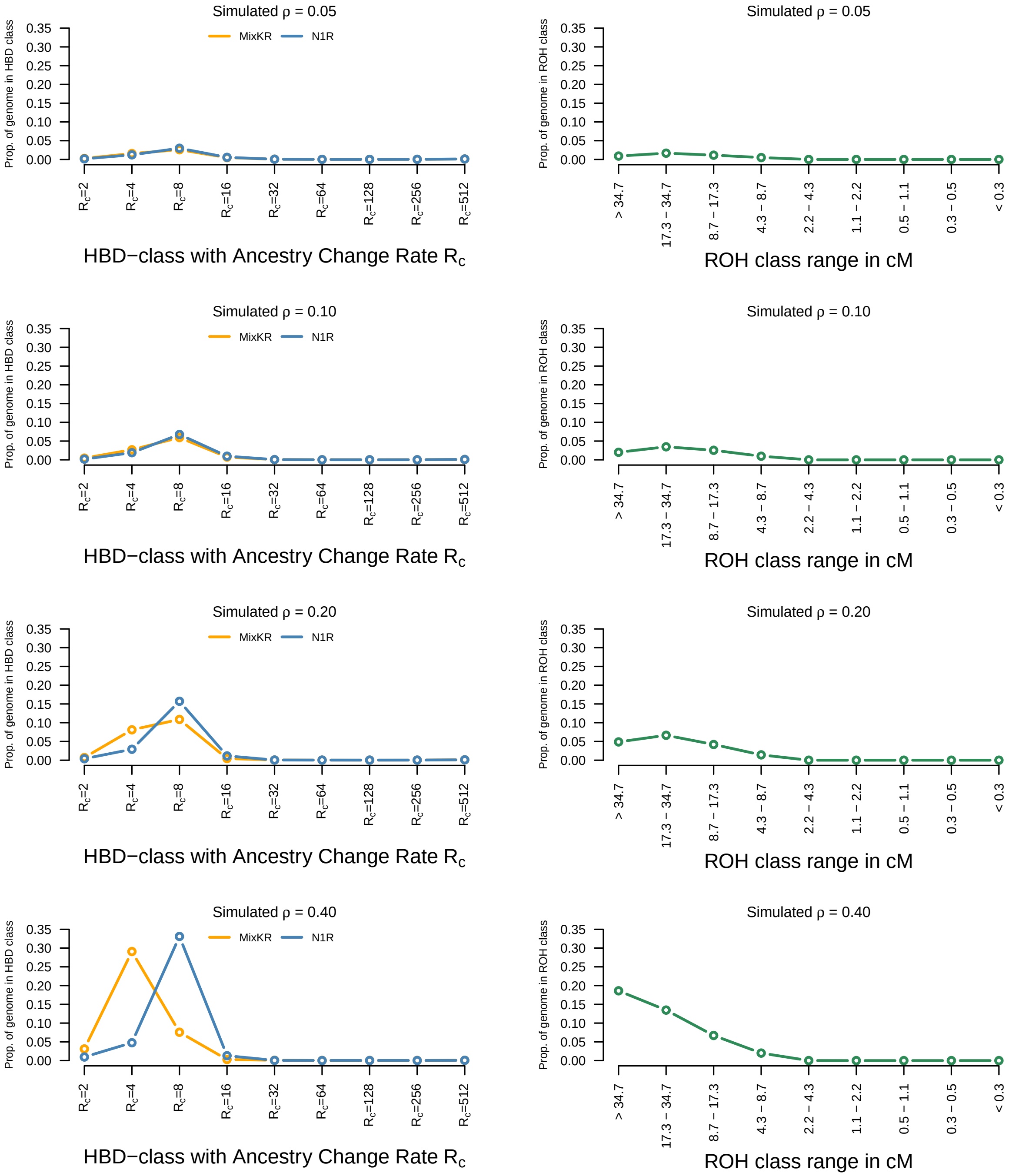


**Supplementary Figure 2. Partitioning in different HBD and ROH classes for data simulated under the 1R model with $R = 8$.** The simulated $\rho$ values are provided above the graphs. The partitioning are realized with the MixKR and N1R models with 9 HBD-classes (left panels). Partitioning in different ROH-classes is performed with GARLIC (right panels), ROH boundaries are selected to match HBD-classes. The boundary between class $c-1$ and $c$ corresponds to the ROH-length for which the exponential distributions with rates $R_{c-1}$ and $R_c$ have equal probability.

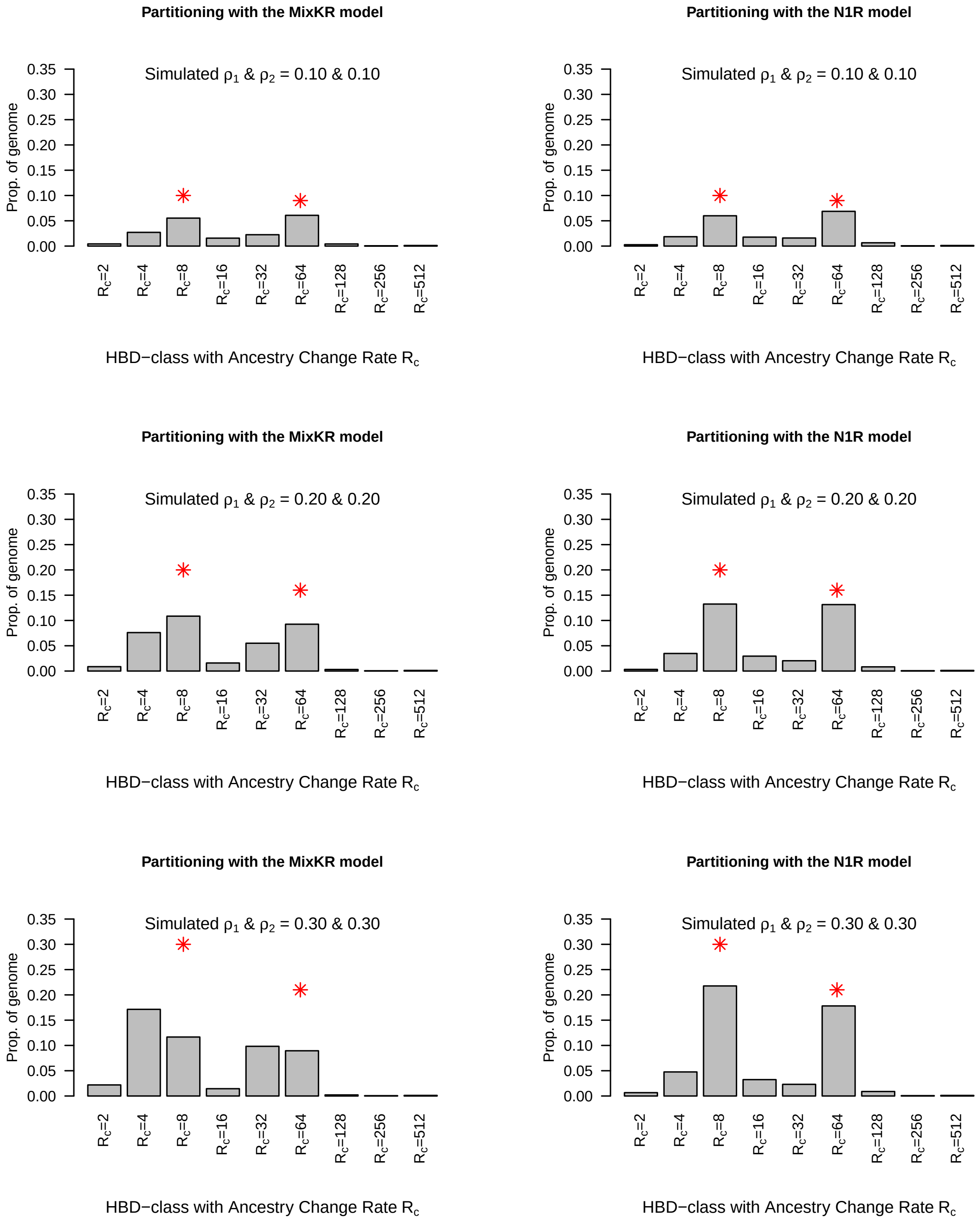


**Supplementary Figure 3. Partitioning in different HBD classes when two HBD classes are simulated under the inference model.** The partitioning are realized with the MixKR (left panels) and N1R (right panels) models. Two HBD classes with $R_c$ equal 8 and 64 are simulated. The simulated $\rho_1$ and $\rho_2$ parameters are indicated above the graphs. The red stars indicate the expected values in different HBD classes, equal to $\rho_1$ and $(1-\rho_1)\rho_2$, respectively.

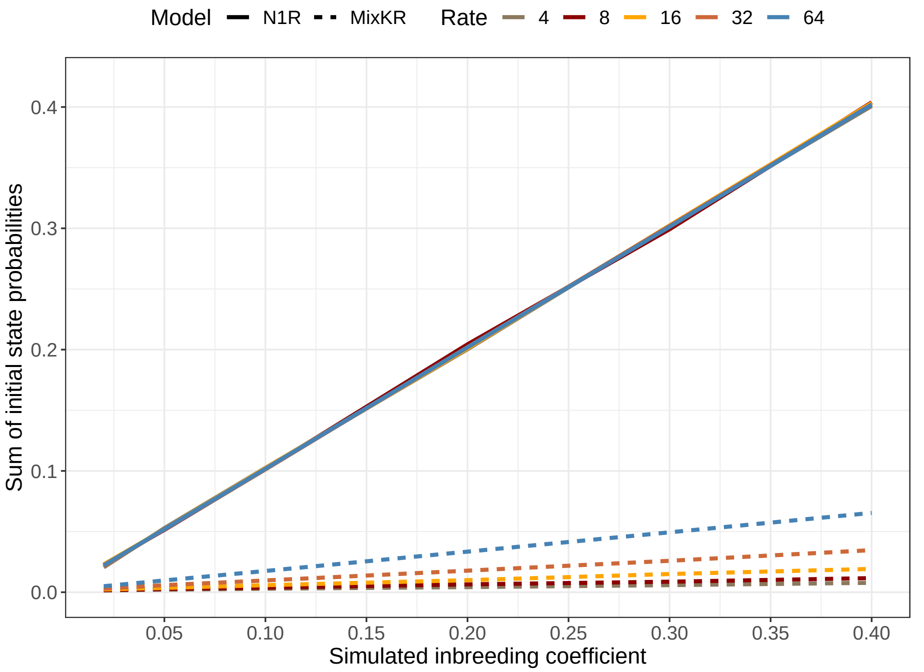


**Supplementary Figure 4. Relationship between initial state probabilities and inbreeding coefficients.** The mixing coefficients $\rho_c$ are used to estimate the initial state probabilities of the HMM with the MixKR and N1R models. For the N1R model, these probabilities are equal to the stationary probabilities and the sum corresponds thus to the inbreeding coefficient $F_\delta$, whereas with the MixKR these values have no interpretation in terms of inbreeding coefficient.

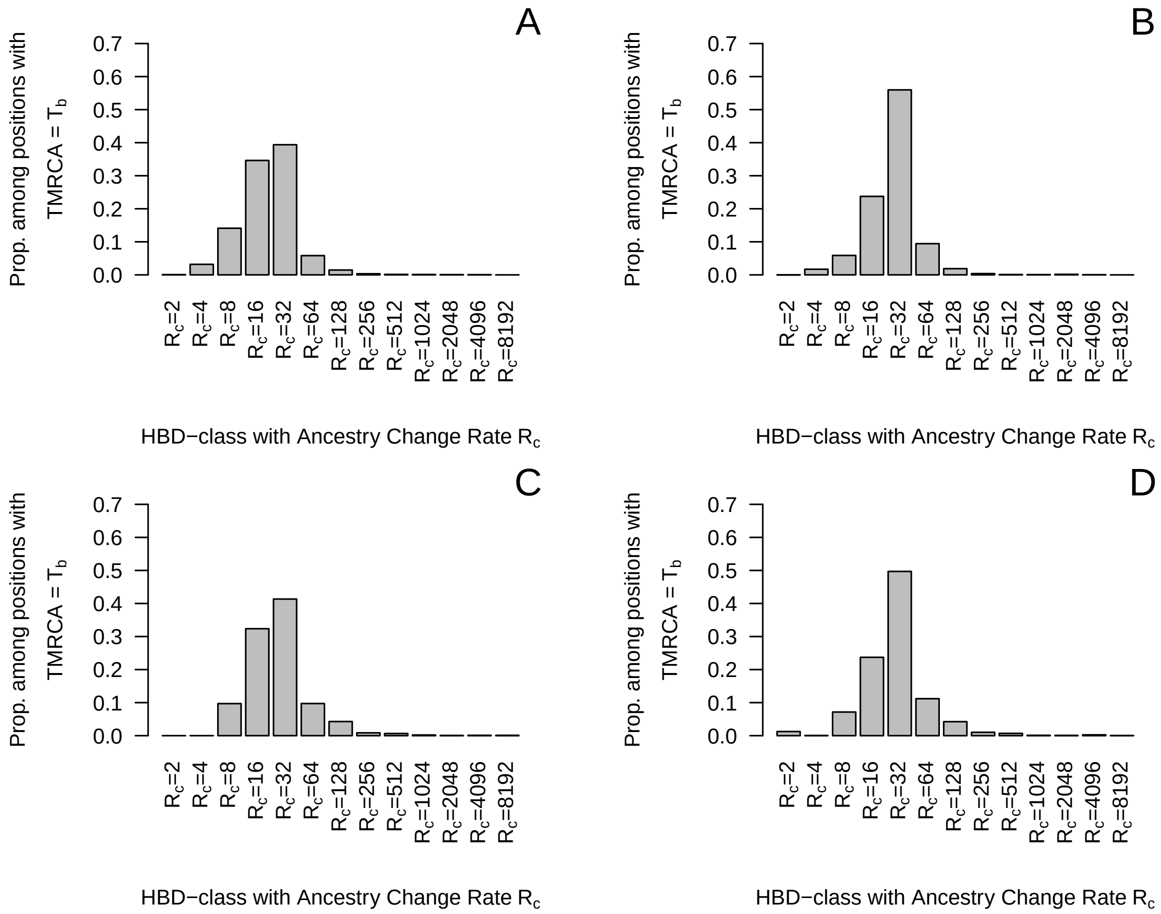


**Supplementary Figure 5. Partitioning of HBD segments related to the bottleneck in different HBD classes.** Data were simulated with a Wright-Fisher process, with a bottleneck in generations 14 to 17 expected to be associated with the HBD class with $R_c = 32$. The simulated $N_e$ during the bottleneck is equal to 20 (A & B) or 50 (C & D). The partitioning is realized with the MixKR (A & C) or N1R (B & D) models.

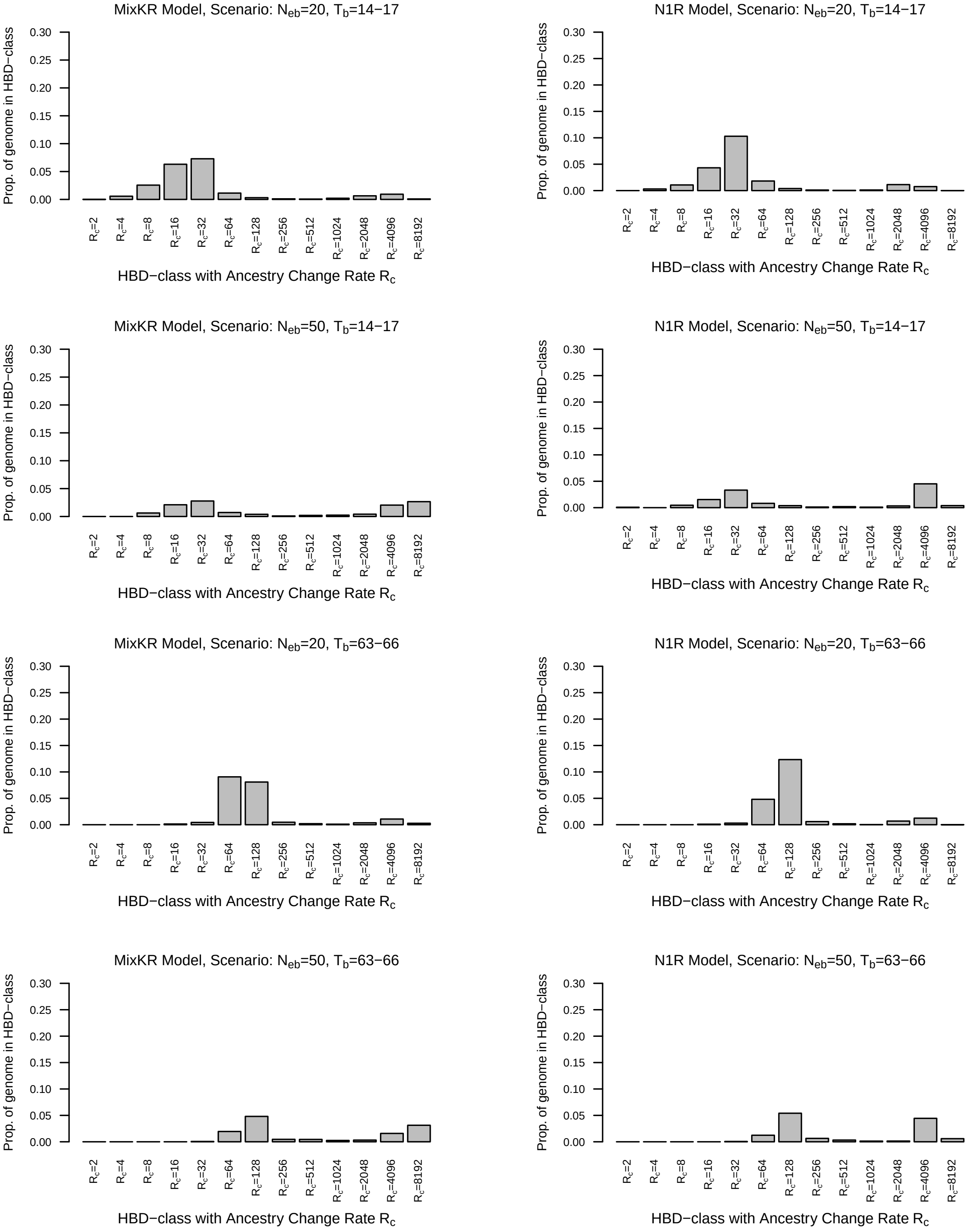


**Supplementary Figure 6. Distributions of the estimated proportion of the individual genomes assigned to each of the 13 predefined HBD classes for the four single bottleneck Wright-Fisher process simulation scenarios.** The partitioning are realized with the MixKR and N1R models. For each scenario, the model and the simulation parameter values are given above the graph (see Material and Methods section in the main text for details).

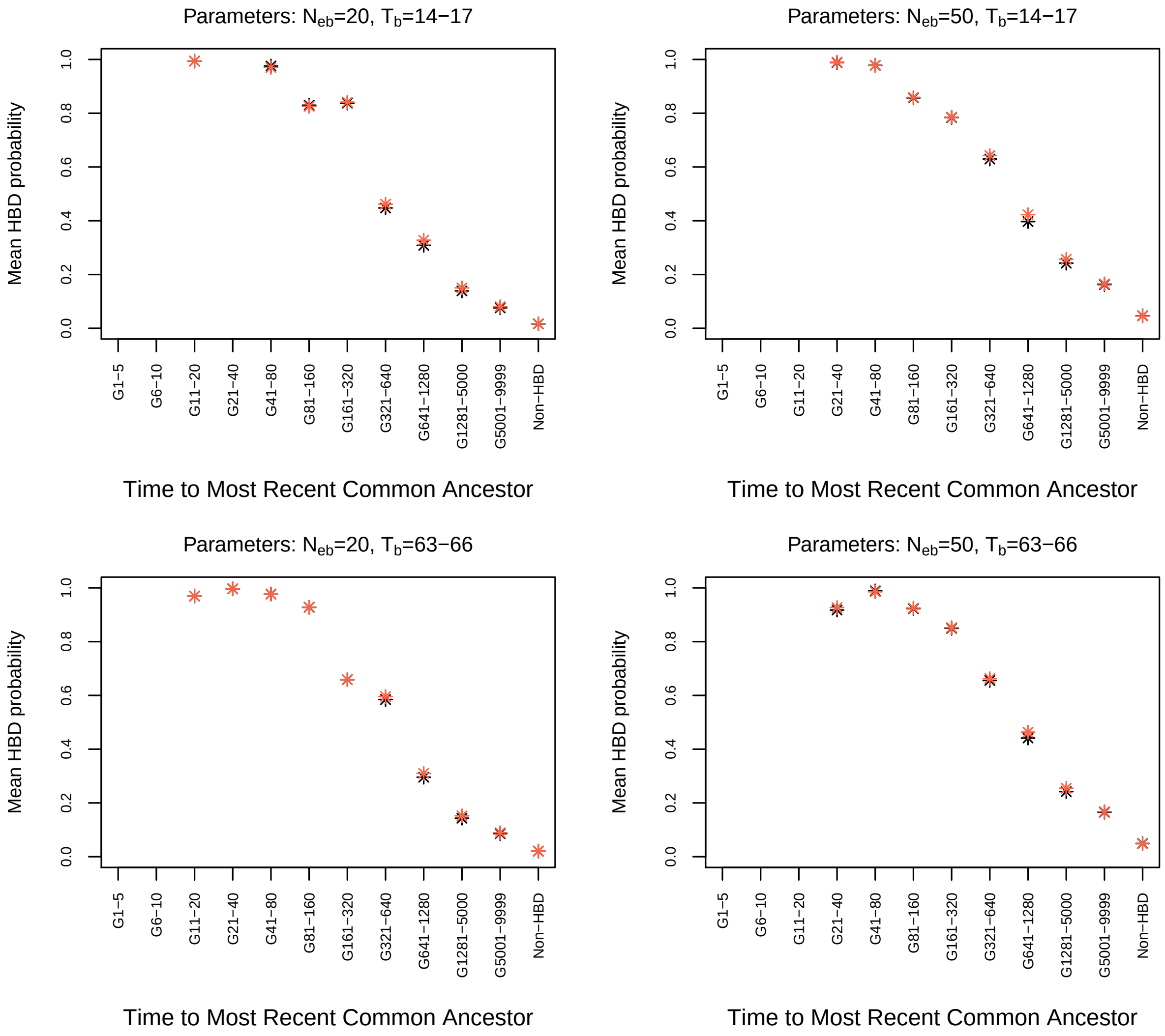


**Supplementary Figure 7. Estimated local inbreeding probabilities ($\phi_m$) averaged over all the simulated individuals and markers as a function of the actual TMRCA of the underlying HBD segments for the four single bottleneck Wright-Fisher process simulation scenarios.** The partitioning are realized with the MixKR (black) and N1R (red) models. For each scenario, the simulation parameter values are given on top of the graph (see Material and Methods section in the main text for details).

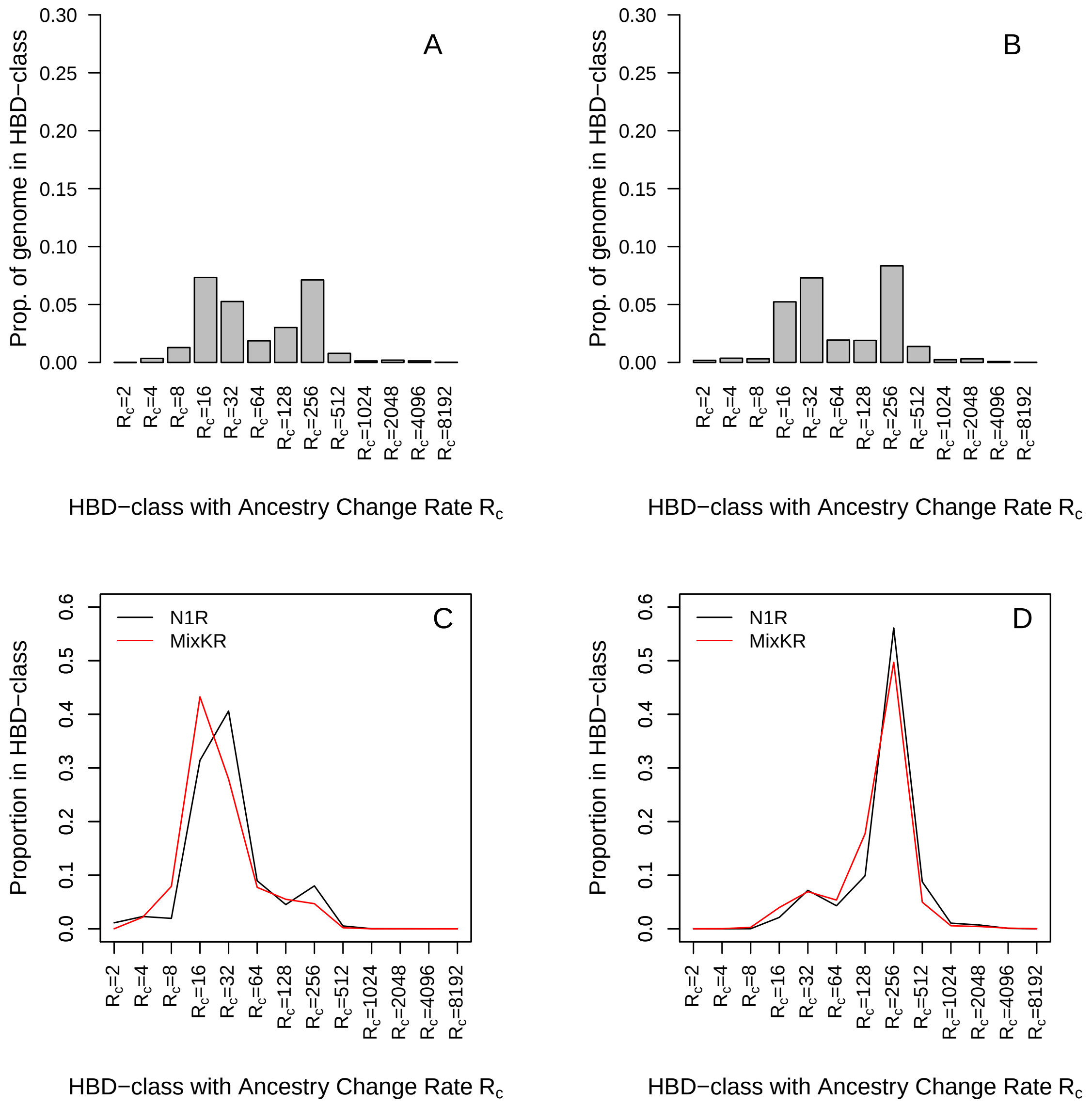


**Supplementary Figure 8. Two bottlenecks scenario ($N_e = 20$).** Data were simulated with a Wright-Fisher process, with two bottlenecks with $N_e = 20$ in generations 14 to 17 and 127 to 130, expected to be associated with the HBD class with $R_c = 32$ and $R_c = 256$. Distributions of the estimated proportion of the individual genomes assigned to each of the 13 predefined HBD classes with A) the MixKR and B) the N1R models. Partitioning of HBD segments related to the two bottlenecks in different HBD classes; C) HBD segments from the first bottleneck (generations 14 to 17) and D) HBD segments from second bottleneck (generations 127 to 130). The partitioning is realized with the MixKR and N1R models.

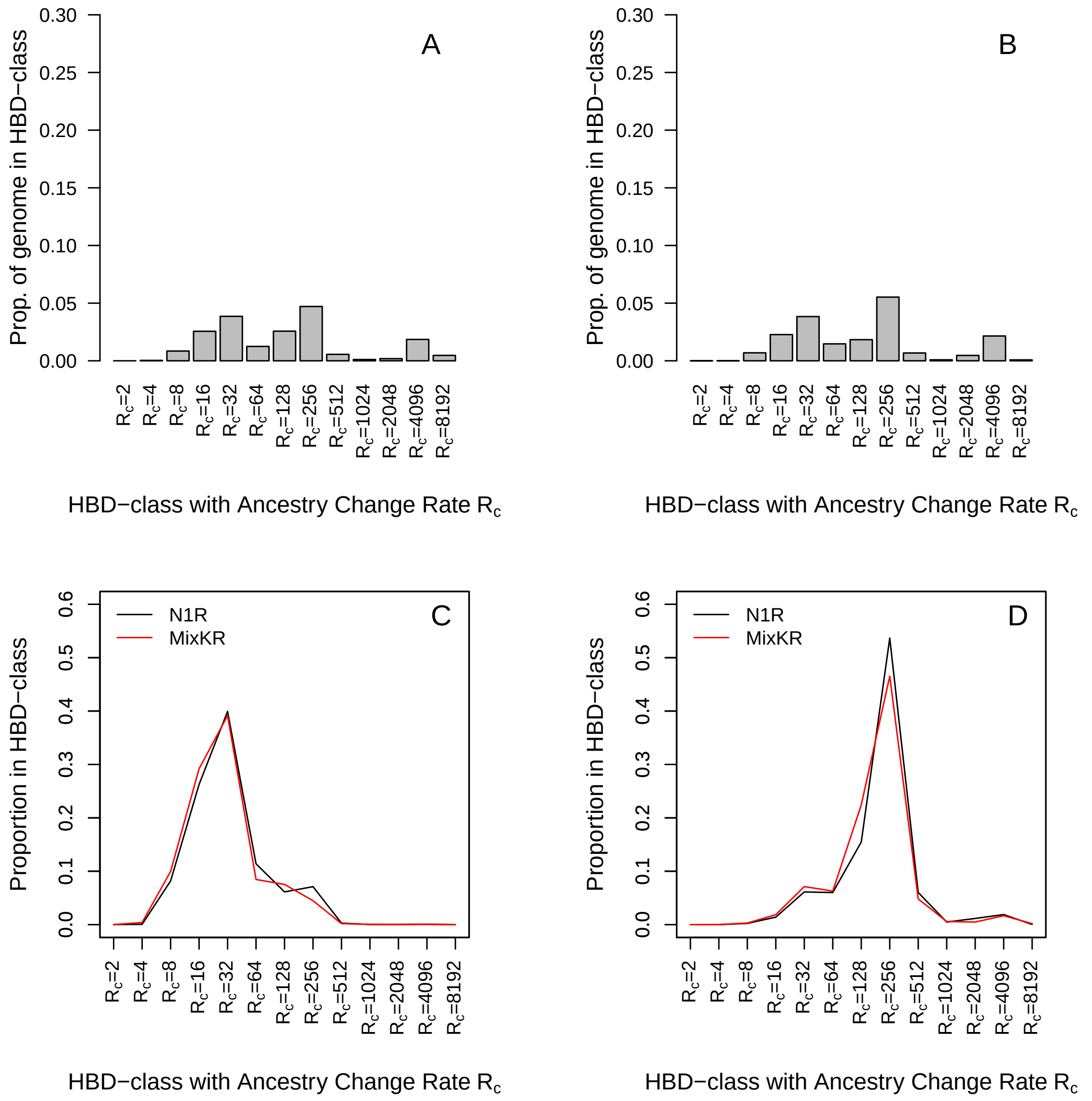


**Supplementary Figure 9. Two bottlenecks scenario ($N_e = 50$).** Data were simulated with a Wright-Fisher process, with two bottlenecks with $N_e = 50$ in generations 14 to 17 and 127 to 130, expected to be associated with the HBD class with $R_c = 32$ and $R_c = 256$. Distributions of the estimated proportion of the individual genomes assigned to each of the 13 predefined HBD classes with A) the MixKR and B) the N1R models. Partitioning of HBD segments related to the two bottlenecks in different HBD classes; C) HBD segments from the first bottleneck (generations 14 to 17) and D) HBD segments from second bottleneck (generations 127 to 130). The partitioning is realized with the MixKR and N1R models.

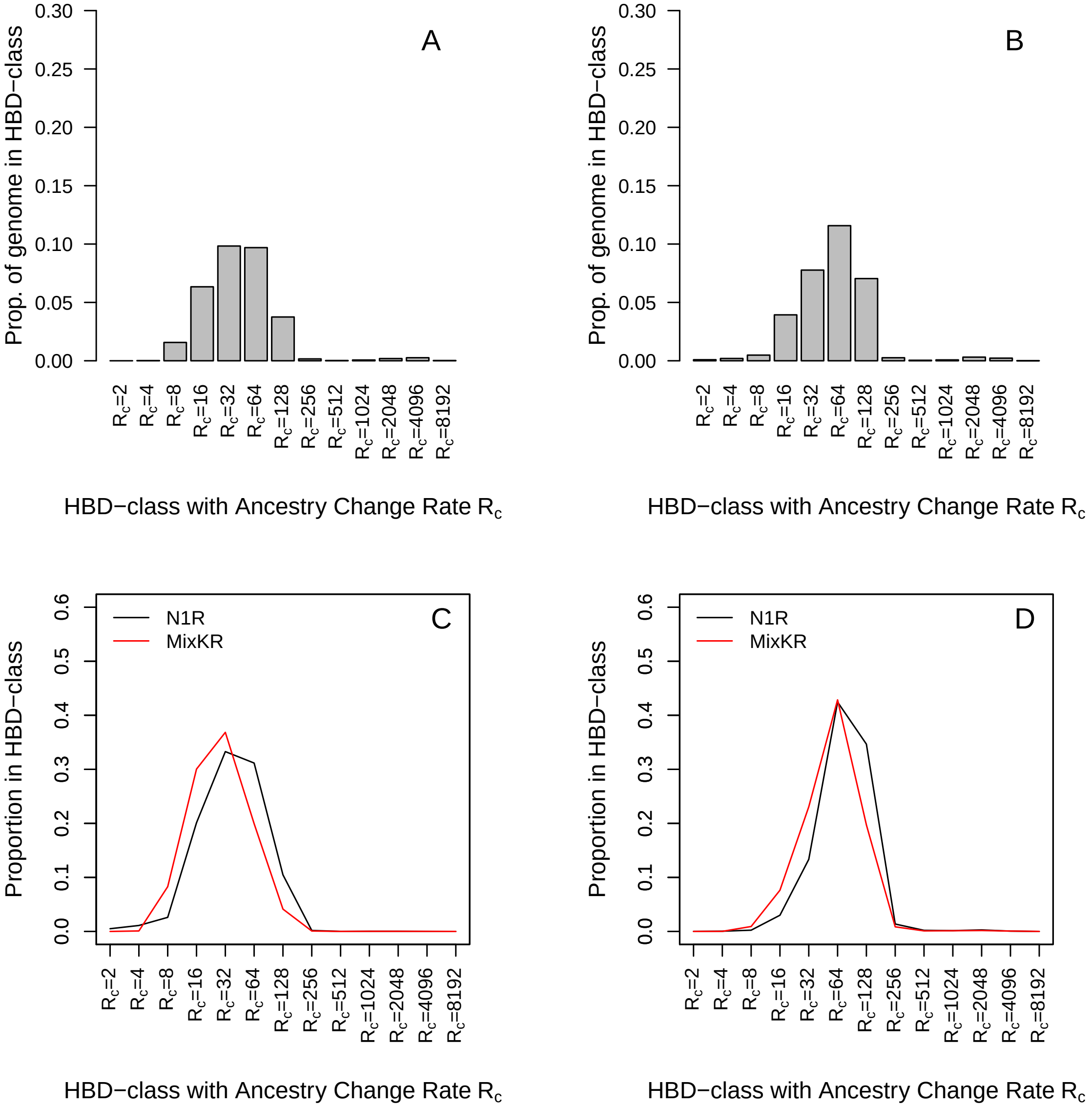


**Supplementary Figure 10. Two bottlenecks scenario ($N_e = 20$).** Data were simulated with a Wright-Fisher process, with two bottlenecks with $N_e = 20$ in generations 14 to 17 and 63 to 66, expected to be associated with the HBD class with $R_c = 32$ and $R_c = 128$. Distributions of the estimated proportion of the individual genomes assigned to each of the 13 predefined HBD classes with A) the MixKR and B) the N1R models. Partitioning of HBD segments related to the two bottlenecks in different HBD classes; C) HBD segments from the first bottleneck (generations 14 to 17) and D) HBD segments from second bottleneck (generations 63 to 66). The partitioning is realized with the MixKR and N1R models.

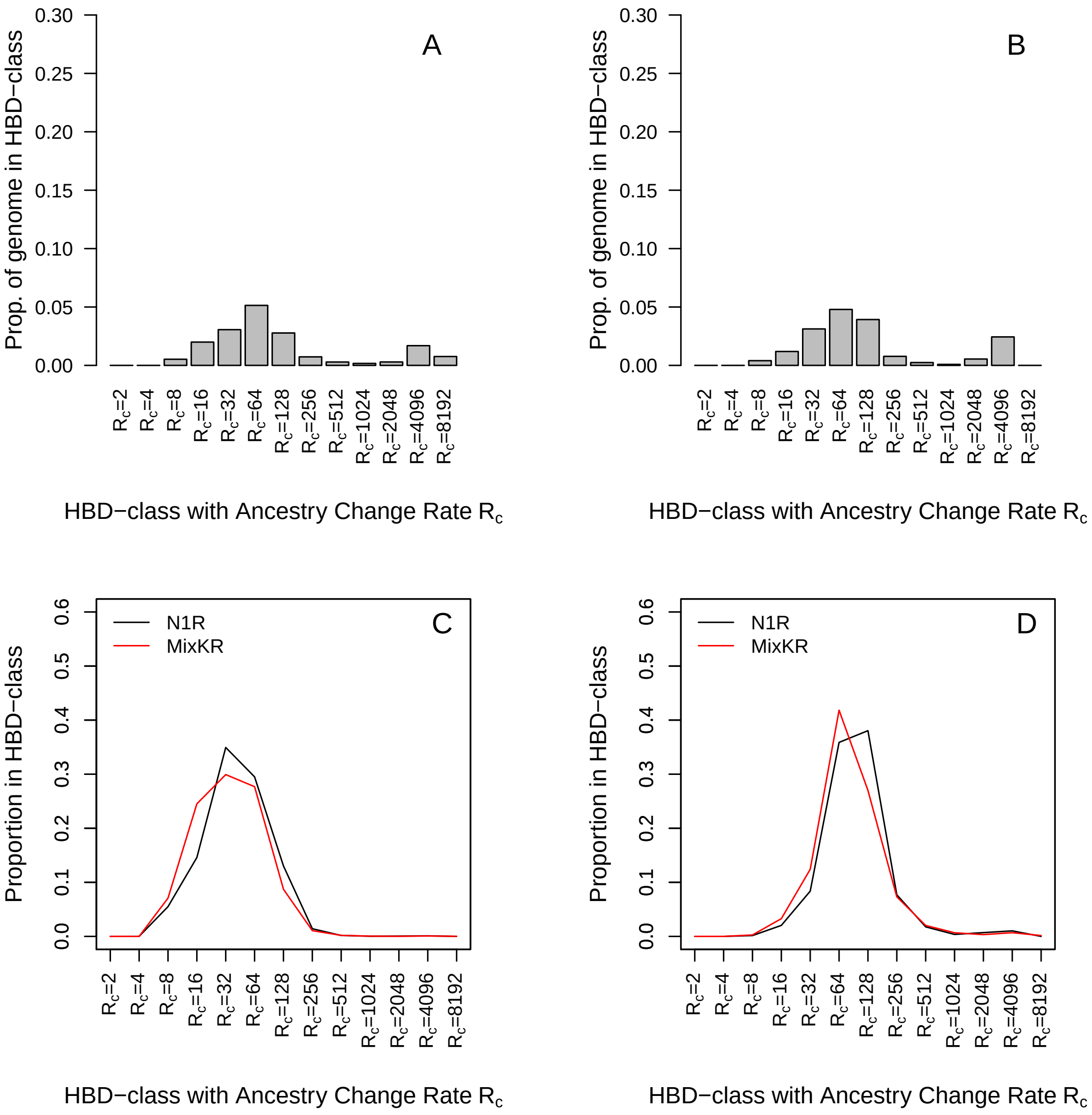


**Supplementary Figure 11. Two bottlenecks scenario ($N_e = 50$).** Data were simulated with a Wright-Fisher process, with two bottlenecks with $N_e = 50$ in generations 14 to 17 and 63 to 66, expected to be associated with the HBD class with $R_c = 32$ and $R_c = 128$. Distributions of the estimated proportion of the individual genomes assigned to each of the 13 predefined HBD classes with A) the MixKR and B) the N1R models. Partitioning of HBD segments related to the two bottlenecks in different HBD classes; C) HBD segments from the first bottleneck (generations 14 to 17) and D) HBD segments from second bottleneck (generations 63 to 66). The partitioning is realized with the MixKR and N1R models.

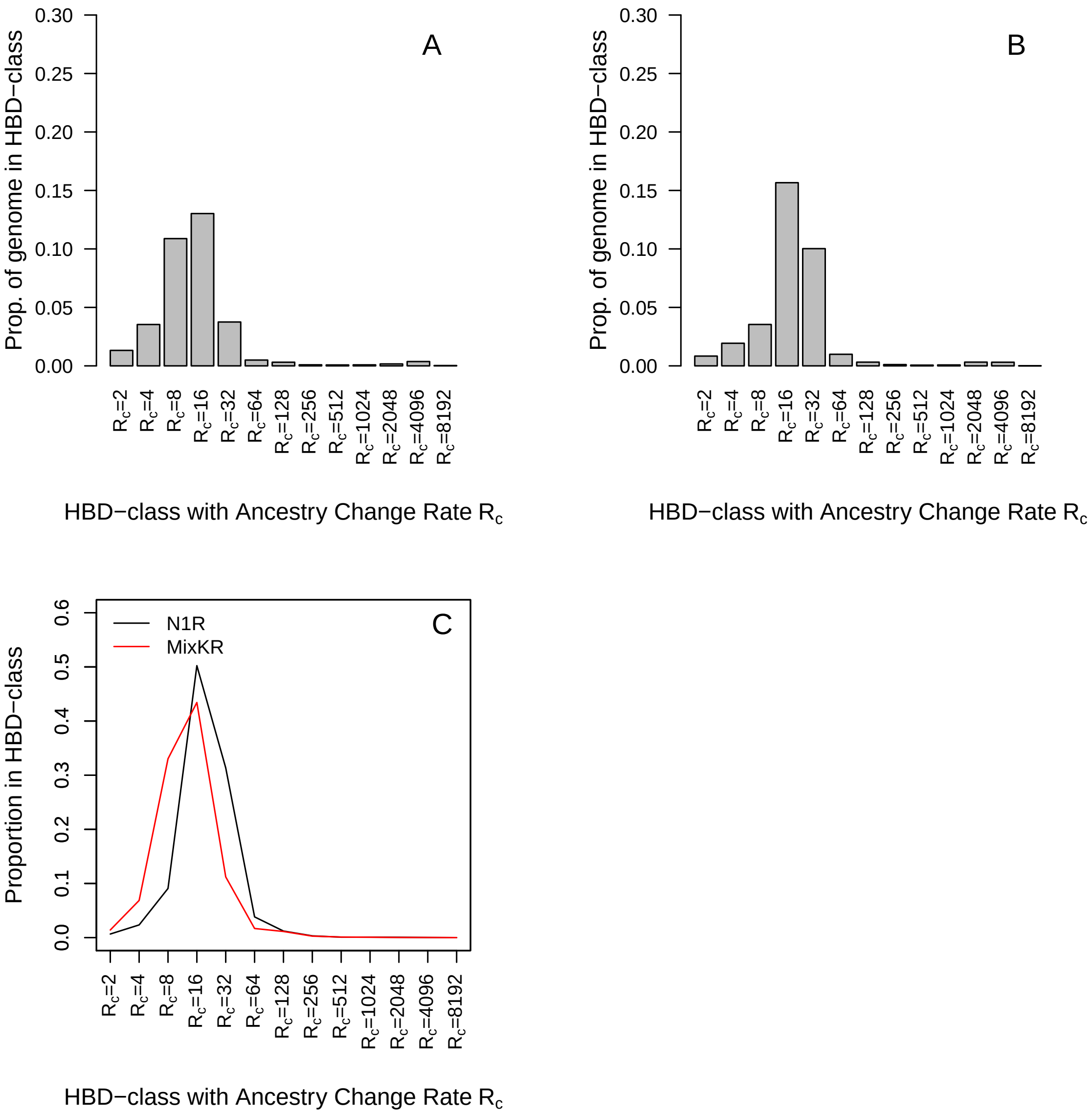


**Supplementary Figure 12. Bottleneck and expansion scenario ($N_e = 20$).** Data were simulated with a Wright-Fisher process, with a bottleneck with $N_e = 20$ in generations 14 to 17, expected to be associated with the HBD class with $R_c = 32$, followed by an expansion with a continuous rate (x 1.19 at each generation) to reach $N_e = 200$ at present. Distributions of the estimated proportion of the individual genomes assigned to each of the 13 predefined HBD classes with A) the MixKR and B) the N1R models. C) Partitioning of HBD segments related to the bottleneck in different HBD classes. The partitioning is realized with the MixKR and N1R models.

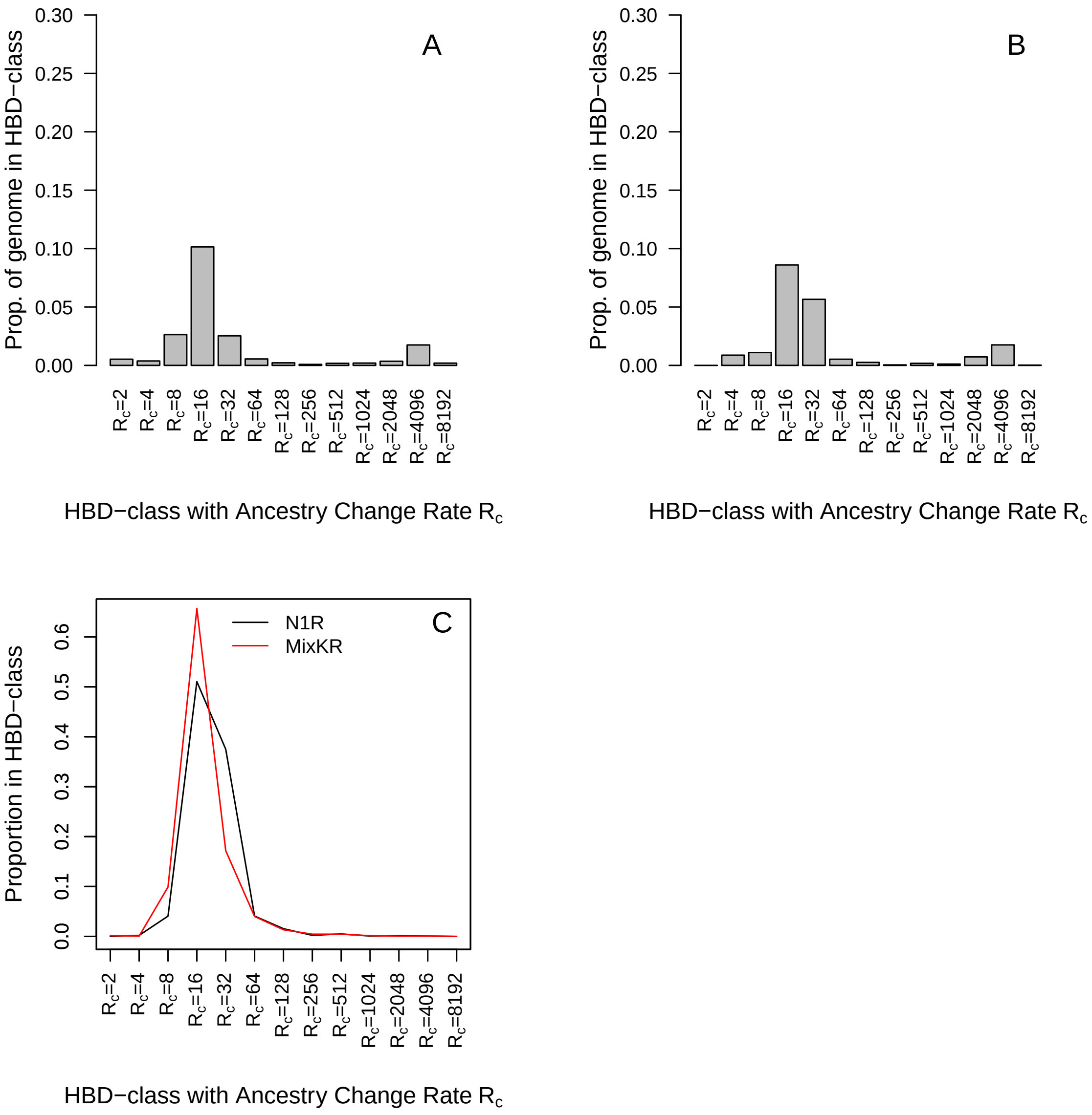


**Supplementary Figure 13. Bottleneck and expansion scenario ($N_e = 50$).** Data were simulated with a Wright-Fisher process, with a bottleneck with $N_e = 50$ in generations 14 to 17, expected to be associated with the HBD class with $R_c = 32$, followed by an expansion with a continuous rate (x 1.19 at each generation) to reach $N_e = 500$ at present. Distributions of the estimated proportion of the individual genomes assigned to each of the 13 predefined HBD classes with A) the MixKR and B) the N1R models. C) Partitioning of HBD segments related to the bottleneck in different HBD classes. The partitioning is realized with the MixKR and N1R models.

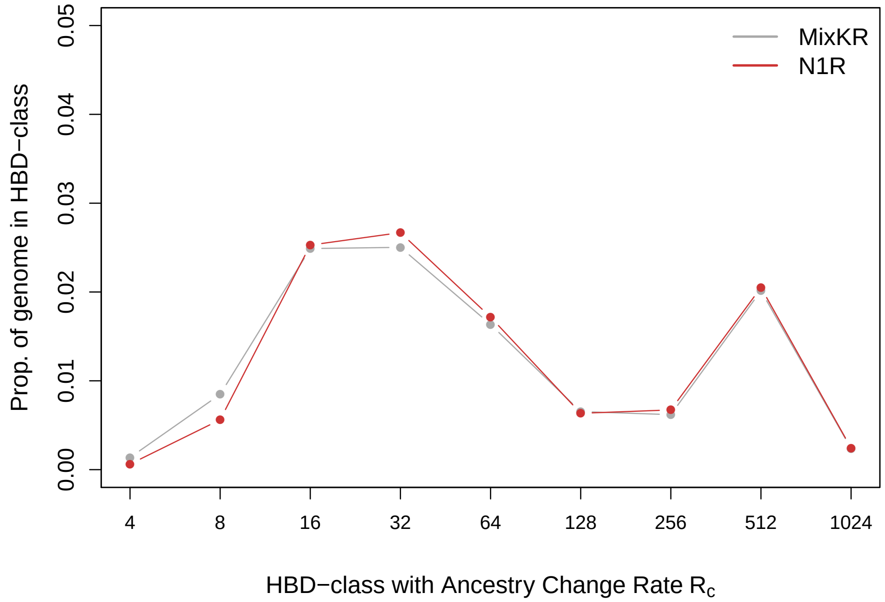


**Supplementary Figure 14. Comparison of HBD partitioning in the Holstein cattle data.** The data consisted in 145 parents genotyped for 37,675 markers (Alemu et al., 2021). The partitioning are realized with the MixKR and the N1R models and with 9 HBD-classes.
